# Pregnancy associated plasma protein-aa (Pappaa) regulates photoreceptor synaptic development to mediate visually guided behavior

**DOI:** 10.1101/257840

**Authors:** Andrew H. Miller, Hollis B. Howe, Bryan M. Krause, Scott A. Friedle, Matthew I. Banks, Brian D. Perkins, Marc A. Wolman

## Abstract

To guide behavior, sensory systems detect the onset and offset of stimuli and process these distinct inputs via parallel pathways. In the retina, this strategy is implemented by splitting neural signals for light onset and offset via synapses connecting photoreceptors to ON and OFF bipolar cells, respectively. It remains poorly understood which molecular cues establish the architecture of this synaptic configuration to split light onset and offset signals. A mutant with reduced synapses between photoreceptors and one bipolar cell type, but not the other, could reveal a critical cue. From this approach, we report a novel synaptic role *pregnancy associated plasma protein aa (pappaa)* in promoting the structure and function of cone synapses that transmit light offset information. Electrophysiological and behavioral analyses indicated *pappaa* mutant zebrafish have dysfunctional cone to OFF bipolar cell synapses and impaired responses to light offset, but intact cone to ON bipolar cell synapses and light onset responses. Ultrastructural analyses of *pappaa* mutant cones showed a lack of presynaptic domains at synapses with OFF bipolar cells. *pappaa* is expressed postsynaptically to the cones during retinal synaptogenesis and encodes a secreted metalloprotease known to stimulate insulin-like growth factor 1 (IGF1) signaling. Induction of dominant negative IGF1 receptor expression during synaptogenesis reduced light offset responses. Conversely, stimulating IGF1 signaling at this time improved *pappaa* mutants’ light offset responses and cone presynaptic structures. Together, our results indicate Pappaa-regulated IGF1 signaling as a novel pathway that establishes how cone synapses convey light offset signals to guide behavior.

**Significance Statement:** Distinct sensory inputs, like stimulus onset and offset, are often split at distinct synapses into parallel circuits for processing. In the retina, photoreceptors and ON and OFF bipolar cells form discrete synapses to split neural signals coding light onset and offset, respectively. The molecular cues that establish this synaptic configuration to specifically convey light onset or offset remain unclear. Our work reveals a novel cue: *pregnancy associated plasma protein aa (pappaa)*, which regulates photoreceptor synaptic structure and function to specifically transmit light offset information. Pappaa is a metalloprotease that stimulates local insulin-like growth factor 1 (IGF1) signaling. IGF1 promotes various aspects of synaptic development and function and is broadly expressed; thus requiring local regulators, like Pappaa, to govern its specificity.

## Introduction

Sensory circuits are wired to translate information in the environment. Many sensory systems detect and code stimuli by their onset and offset. Neural signals conveying ON/OFF input are split into parallel circuits for processing (Chalasani et al., 2007a; Scholl et al., 2010; Gallio et al., 2011; Gjorgjieva et al., 2014). This split occurs at discrete circuit loci, like synapses between sensory neurons and interneurons (DeVries et al., 2006; Westheimer, 2007). Synapses epitomize the notion that form follows function, and thus their architecture mediating signal splitting must be established during development. For some circuits, the synapses that split ON/OFF signals have been identified and physiologically characterized (Wassle, 2004), yet the molecular cues that establish these synapses remain poorly understood. Defining these cues and their contribution to synaptic architecture will inform how synapses are built to split ON/OFF signals.

Visual circuits exemplify the splitting of ON/OFF stimuli. Retinal photoreceptor neurons respond to light onset and offset by modulating their release of glutamate, which differentially affects the activity of discrete postsynaptic bipolar cell populations (Wassle, 2004; Hoon et al., 2014). A cone photoreceptor’s terminal establishes discrete synapses with invaginating dendrites of both ON and OFF bipolar cells to convey information about light onset and offset, respectively (Fig. 1A–B). At the cone terminal’s center, ON bipolar cell dendrites express metabotropic glutamate receptors and are flanked by horizontal cell dendrites (Haverkamp et al., 2000). Apposed to ON bipolar cell dendrites, the cone docks synaptic ribbons to mediate synaptic vesicle release (Schmitz, 2014). OFF bipolar cell dendrites invaginate peripheral regions of a cone’s terminal and form “flat contacts” with the cone (Nelson and Connaughton, 1995). Flat contacts are identifiable by electron rich pre-and postsynaptic densities (Dowling and Boycott, 1966). The postsynaptic domain includes clustered ionotropic, AMPA-type glutamate receptors on the OFF bipolar cell dendrites, whereas the presynaptic domain’s composition remains unknown (Boycott and Hopkins, 1993; Tsukamoto and Omi, 2015).

**Figure 1.**
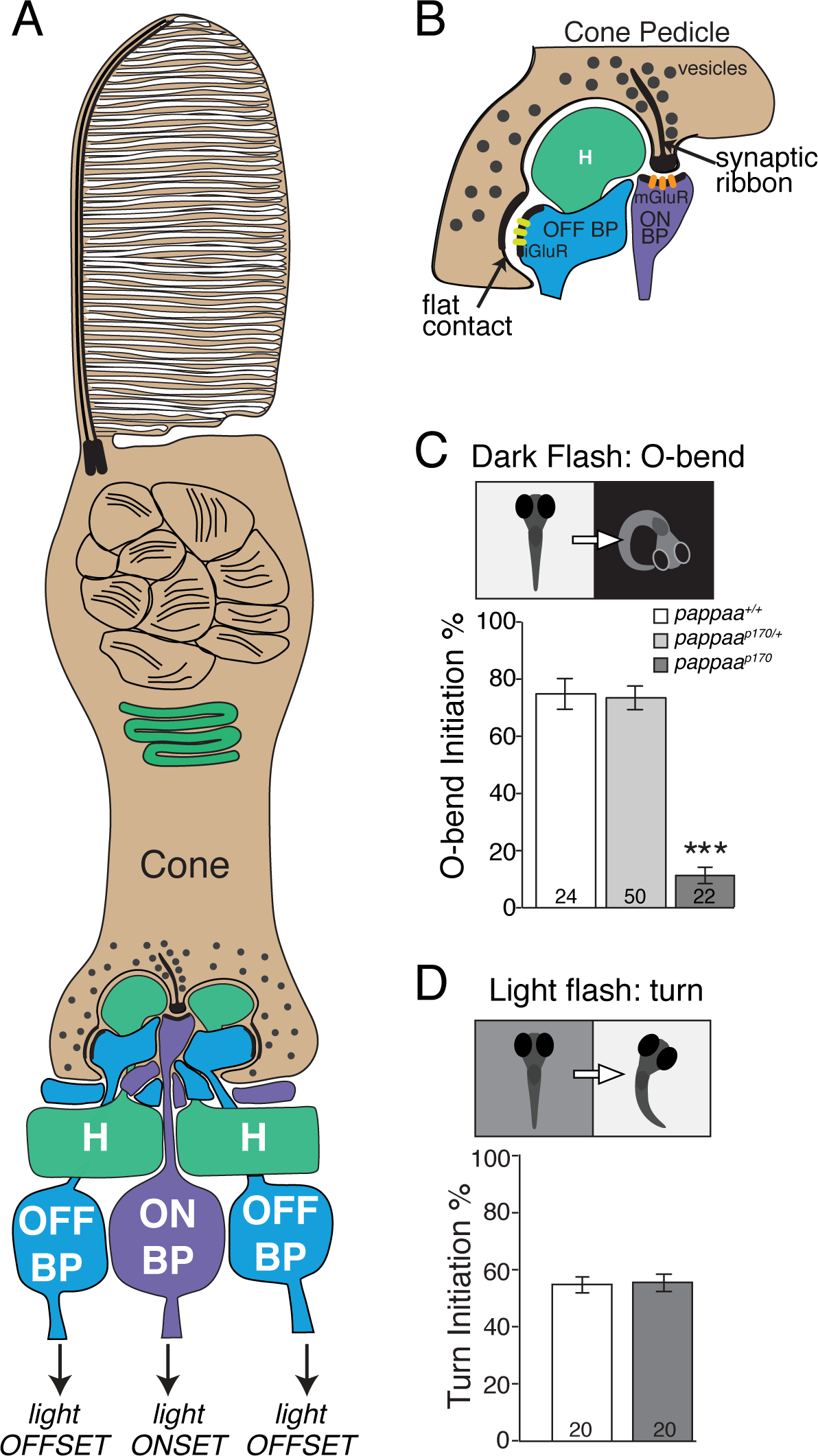
*pappaa*^*p170*^ larvae show impaired initiation of behaviors to light offset. (A) Cartoon of a cone photoreceptor and its synaptic connections with dendrites of ON (purple) and OFF bipolar cells (blue) and horizontal cells (H, green) within the retinal outer plexiform layer. (B) Higher magnification view of contacts between cone and ON and OFF bipolar cells that indicates differences in synaptic structure. At cone to ON bipolar cell contacts, a synaptic ribbon is docked presynaptically and metabotropic glutamate receptors (mGluR) cluster at the postsynaptic density. At cone to OFF bipolar cell contacts, flat contacts are marked by pre-and postsynaptic densities and AMPA-type ionotropic glutamate receptors (iGluR) clusters postsynaptically. (C) Dark flash stimulus elicits O-bend response. Mean frequency of O-bend responses initiated to a series of 10 dark flash stimuli. ***p<0.001, ANOVA with Bonferroni correction. (D) Light flash elicits a low angle turn response. Mean frequency of turns initiated to a series of 10 light flash stimuli. N at base of bars = number of larvae. Error bars indicate SEM.

Which molecular cues establish this synaptic configuration to specifically mediate the transmission of light onset or offset? A powerful tool to identify a cue is a mutant with dysfunctional synapses between photoreceptors and one bipolar cell type, but not the other. Zebrafish are ideal to isolate such a mutant, given the model’s amenability to forward genetic screens for visual defects and their retinal circuits’ rapid development and homology to mammals (Brockerhoff et al., 1995; Easter and Nicola, 1996; Neuhauss et al., 1999; Fadool and Dowling, 2008). By 5 days of age cone to bipolar cell synapses exhibit mature physiology and drive stereotyped behavioral responses to light onset and offset (Biehlmaier et al., 2003; Fleisch and Neuhauss, 2006; Burgess and Granato, 2007b; Burgess et al., 2010). Indeed, zebrafish mutants have revealed genes underlying the formation of photoreceptors’ synapses to both bipolar cell types and photoreceptors’ synapses with ON bipolar cells (Brockerhoff et al., 1995; Neuhauss et al., 1999; Allwardt et al., 2001). These genes have been shown to regulate phototransduction, cell survival, vesicle recycling, calcium homeostasis, and cytoskeletal dynamics (Stearns et al., 2007; Lewis et al., 2011; Jia et al., 2014; Wasfy et al., 2014; Daniele et al., 2016; Lin et al., 2016; Shi et al., 2017). However, mutants have yet to reveal a cue that specifically regulates the formation of synapses between photoreceptors and OFF bipolar cells; and thus, specifically promotes transmission of light offset signals. We identified a zebrafish mutant that fills this void based on analyses of behavior, synaptic structure, and physiology. These mutants harbor nonsense mutations in *pregnancy associated plasma protein aa (pappaa)*, which encodes a secreted metalloprotease that stimulates local insulin-like growth factor (IGF) signaling (Lawrence et al., 1999; Conover, 2012; Oxvig, 2015). A role for Pappaa in synaptic development and function is novel. Here, we describe how Pappaa-IGF1R signaling influences cone to OFF bipolar cell synaptic structure and function and behavioral responses to light offset.

## Materials and Methods

### Fish Maintenance and breeding

Larval zebrafish *(Danio rerio)* were raised from crosses of the following adults: Tubingen long-fin wild type; *pappaa^p170/+^; Tg(isl2b-gfp); Tg(hsp70:dnlGFlRa-EGFP); pappaa^p170/+^, Tg(isl2b-gfp);* or *pappaa*^*p170U*^, *Tg(hsp70:dnIGF1Ra-GFP).* All lines were maintained on a Tubingen long-fin wild type background. From fertilization, embryos and larvae were raised at 29°C in E3 medium (5 mM NaCl, 0.17 mM KCl, 0.33 mM CaCl_2_, 0.33 mM MgSO_4_, pH adjusted to 6.8-6.9 with NaHCO_3_) on a 14:10 hour light:dark cycle. To inhibit melanization, larvae used for fluorescent microscopy were raised in phenylthiourea (PTU, 0.2mM in E3 medium) in the dark beginning at 16-22 hours post-fertilization (hpf). All experiments were conducted between 4-10 days post-fertilization (dpf). Larvae raised beyond 6 dpf were fed paramecia. Genotyping of the *pappaa*^*p170*^ allele was performed as previously described (Wolman et al., 2015).

### Behavioral Assays, Video Recording, and Analysis

Behavioral responses to light offset (dark flash), light onset (light flash), light adaptation, and phototaxis stimuli were recorded using a MotionPro Y4 high-speed video camera (Integrated Design Tools, Tallahassee, FL, USA) at 1,000 frames per second (unless otherwise noted) and 512×512 pixel resolution, using a 50mm macro lens (Sigma Corporation of America, Ronkonkoma, NY, USA). Larvae were illuminated from above with a mounted LED light (MCWHL5 6500 K LED, powered by LEDD1B driver, Thorlabs, Newton, New Jersey, USA) and below with an infrared light source (IR Illuminator CM-IR200B, C&M Vision Technologies, Houston, TX, USA). Video images were analyzed for turn and swim initiations and kinematics with the FLOTE software package as previously described (Burgess and Granato, 2007b, a; Burgess et al., 2010; Wolman et al., 2011). On the day of testing, the 5 dpf larvae were held in 60mm Petri dishes with 20 larvae in 7mL E3 and were adapted on a white light box (800 μW/cm^2^) for at least 1 hour prior to testing as described below.

Light offset-induced (dark flash) O-bends and light onset-induced (light flash) turns were elicited and analyzed as previously described (Burgess and Granato, 2007b; Wolman et al., 2011). For data represented in Fig. 3C, larvae were tested in 60mm Petri dishes at the density indicated above. For data shown in Fig. 12A, *pappaa*^*p170*^ larvae were identified on 4 dpf based on their failure to habituate to acoustic stimuli (Wolman et al., 2015) and then grouped in 60mm Petri dishes with 20 larvae and 7mL E3 in each dish for testing. For data shown in Fig. 1C-D, 3B, and 3D-H the larvae were transferred from Petri dishes to a 4×4 grid for testing, and then genotyped for*pappaa*^*p170*^ post-hoc (Wolman et al., 2011; Wolman et al., 2015). In the dish or grid, larvae were acclimated to the illuminated testing stage (85 μW/cm^2^ for dark flashes, 25 μW/cm^2^ for light flashes) for 5 minutes before exposure to a series of 10 flash stimuli (dark flash = lights off, light flash = light increased to 600 μW/cm^2^ within 1msec). Each stimulus lasted for 1 second and stimuli were presented at a 30 second interstimulus interval.

Light adaptation was performed as previously described (Burgess and Granato, 2007b). *pappaa*^*p170*^ larvae were identified on 4 dpf based on their failure to habituate to acoustic stimuli (Wolman et al., 2015) and then grouped in 60mm Petri dishes with 20 larvae and 7mL E3 in each dish. Larvae were acclimated on the testing stage for 1 hour in either darkness (dark to light group, DL) or light (85 μW/cm^2^, constant light group, CL). For the DL group, light onset is marked at the 0 timepoint. The CL group remained in constant light (85 μW/cm^2^) at the 0 timepoint. To determine turn and swim initiations, 500ms long video recordings were captured at each time point indicated in Fig. 2A-B.

**Figure 2.**
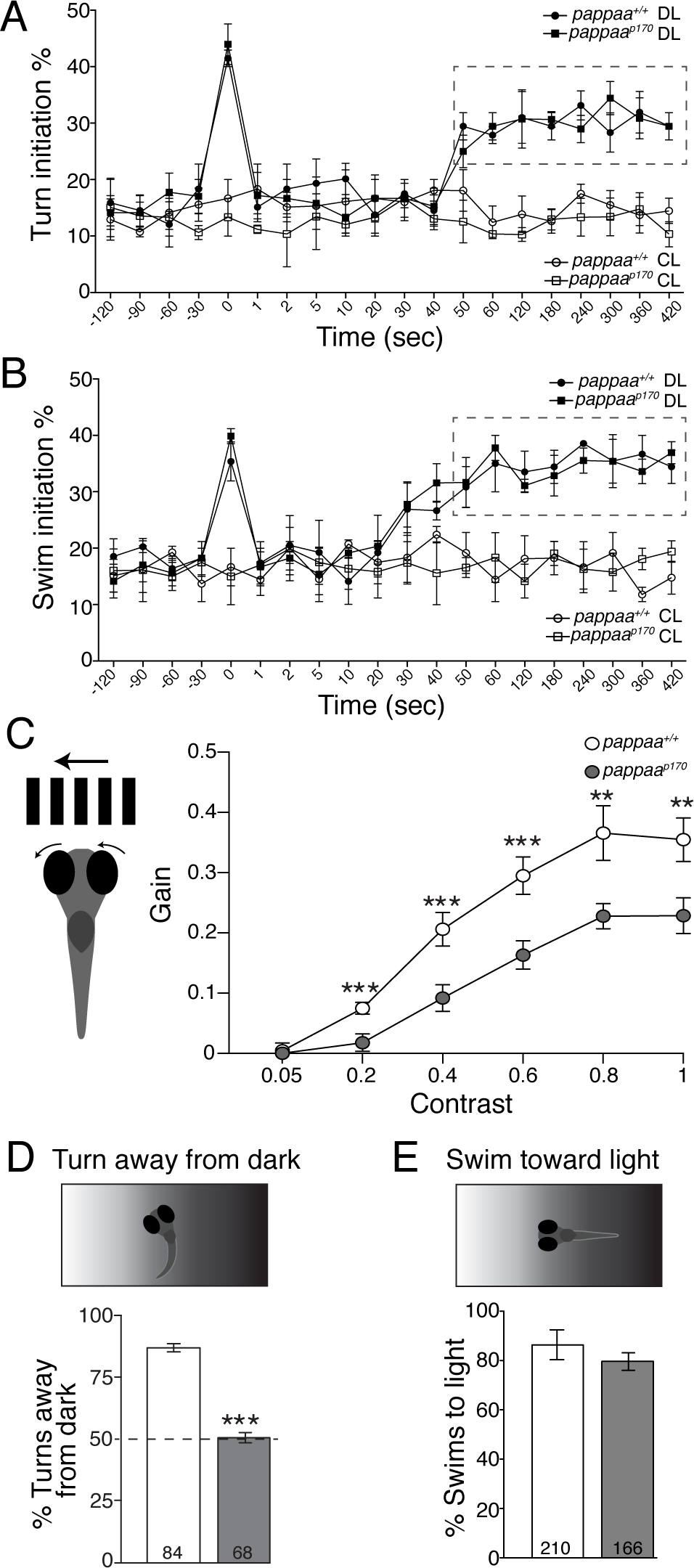
Light adaptation and contrast-mediated behavior by *pappaa*^*p170*^ larvae. (A-B) Mean frequency of turn (A) or swim (B) bout initiation per 1s interval beginning at time on X-axis. DL indicates larvae transitioned from dark to light at time = 0. CL indicates larvae were in constant light throughout assay. Dashed box marks the period in which DL larvae initiated more turns and swims compared to CL larvae of the same genotype, indicative of light adaptation. p<0.001, T-test between like genotypes at each time point. N = 3 groups of 20 larvae/group. (C) Optokinetic reflex (OKR) allows larval eyes to track movement of alternating bars of varying contrast. Mean OKR gain (eye velocity/stimulus velocity) at varying contrasts. N= 8 larvae per genotype. **p<0.01, ***p<0.001, T-tests between genotypes at each contrast value. (D-E) To perform phototaxis, larvae positioned perpendicular to a light gradient stereotypically turn away from darkness (D) and once positioned toward the light, a larva will initiate swim bouts (E). (D) Mean percentage of turns initiated away from darkness by larvae positioned between 75-105 degrees to the light-dark gradient. ***p<0.001, T-tests between genotypes. (E) Mean frequency of swim bouts initiated by larvae facing +/− 30 degrees of the light stimulus. N at base of bars = number of trials in which a turn (D) or swim (E) was evaluated based on larval position with respect to target light. Error bars indicate SEM.

Phototaxis was performed as previously described (Burgess et al., 2010). *pappaa*^*p170*^ larvae were identified at 4 dpf based on their failure to habituate to acoustic stimuli (Wolman et al., 2015) and then grouped in 60mm Petri dishes with 20 larvae and 7mL E3 in each dish for testing. Each dish of larvae was acclimated to an illuminated testing stage (25 μW/cm^2^) for 5 minutes. To elicit phototaxis, the uniform illumination was removed, revealing a small target light (10 μW/cm^2^) on one side of the arena for 30 seconds. Larvae initially positioned on the side (half) of the dish with the target light were excluded from analysis. Larvae oriented perpendicular to the target light (75° - 105°) were analyzed for turn directionality (Fig. 2D), and larvae facing the target light (-30° - 30°) were analyzed for swims toward the target (Fig. 2E). Each dish was tested for phototaxis 3 times, with 5 minutes of uniform illumination (25 μW/cm^2^) between trials.

Optokinetic response (OKR) measurements were performed as previously described (Daniele et al., 2016; Lessieur et al., 2017). Larvae at 5 dpf were assessed for contrast sensitivity using a VisioTracker system (VisioTracker 302060 Series, TSE Systems, GmbH Bad Homburg, Gemany). A stimulus was presented for 3 seconds before reversing direction for another 3 seconds and stimuli were presented with a constant angular velocity of 7.5 degrees per second. Genotyping was performed post-hoc.

### Pharmacology

Larvae were treated with the following compounds by addition of the compound to the larvae’s E3 media: SC-79 (Tocris), recombinant IGF1 (Cell Sciences), and AMPA (Tocris). SC-79 was dissolved in 100% DMSO and administered in a final concentration in 1% DMSO in E3. IGF1 was dissolved in 10mM HCL to 1mg/mL and further diluted in E3 to be administered at a desired final concentration. AMPA was dissolved and administered in E3. Control treatments were either 1% DMSO in E3 (for SC-79) or E3 only (for IGF1, AMPA). For treatments over multiple days, the compound/E3 was replaced every 12 hours. For acute treatments on 5 dpf, larvae were bathed in the compound/E3 for 20 minutes prior to and during the behavioral testing. Doses of each compound were prescreened for potential effects on embryonic and larval health, gross behavior, baseline startle responsiveness and the stereotyped kinematic parameters of O-bend behavior (Burgess and Granato, 2007b).

### *Pappa* and *dominant negative IGF1Ra* misexpression

For human *pappa* RNA injection, full length human *pappa* and *pappa^E483A^* constructs (Boldt et al., 2001) were transcribed using the mMessage mMachine kit (Ambion) and injected at the 1-cell stage at doses ranging from 1-200 picograms. Embryos injected with greater than 100pg *pappa or pappa^E483A^* mRNA showed gross morphological abnormalities and necrosis, whereas embryos injected with 100pg *pappa* (or less) appeared morphologically normal. 100pg injections were used to generate data in Fig. 3B. *pappa* mRNA expression at 5 dpf was validated as previously described (Wolman et al., 2015).

**Figure 3.**
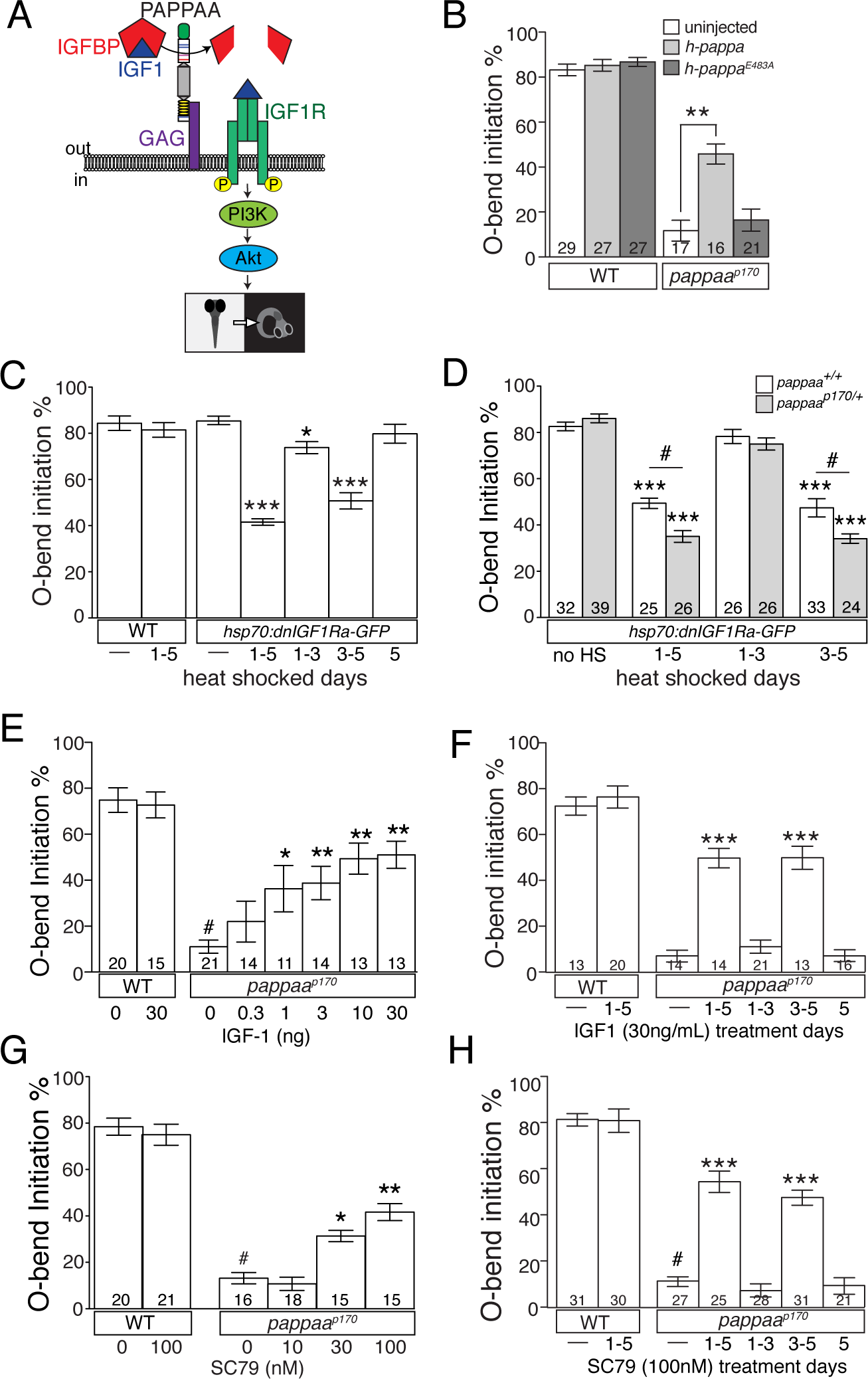
PAPPAA-IGF1R signaling promotes O-bend initiation to light offset. (A) Schematic of Pappaa regulated IGF1 signaling. Pappaa is secreted, binds cell surface glycosaminoglycan (GAG) and cleaves IGF binding proteins (IGFBP) to increase bioavailability of IGF1. “Free” IGF1 binds and activates the IGF1R, which stimulates PI3K and Akt to promote O-bend initiation at light offset. (B) Mean initiation frequency of O-bend responses at 5 dpf after injection of human wild type *pappa* mRNA or a proteolytically inactive *pappa^E483A^* mRNA. **p<0.01 versus uninjected, ANOVA with Bonferroni correction. (C-H) Mean initiation frequency of O-bend responses at 5 dpf in the following conditions/experimental groups: (C) Heat shock or no heat shock of wild type or *Tg(hsp70:dnlGF1Ra-GFP)* larvae. *p<0.05, ***p<0.001 vs. no heat shock *Tg(hsp70:dnlGF1Ra-GFP)* larvae, ANOVA with Bonferroni correction. N= 5 groups of 15 larvae per treatment condition. (D) Heat shock or no heat shock of *Tg(hsp70:dnlGF1Ra-GFP)* larvae in *pappaa*^*+,+*^ or *pappaa*^*p170U*^ background. ***p<0.001 versus no heat shock *Tg(hsp70:dnlGF1Ra-GFP)* of same genotype, ANOVA with Bonferroni correction. #p<0.05, T-test. N larvae shown at base of bars. (E-H) Treatment with recombinant IGF-1 (E-F) or SC79 (G-H) from 3-5 dpf (E,G) or the days indicated (F,H). #p<0.001 vs. untreated wild type, *p<0.05, **p<0.01, ***p<0.001 vs. untreated *pappaa*^*p170*^. ANOVA with Bonferroni correction. N larvae shown at base of bars. All error bars indicate SEM.

To induce expression of dominant negative *IGF1Ra-EGFP*, we heat shocked *Tg(hsp70:dnlGF1Ra-EGFP)* larvae for 1 hour every 12 hours at 37°C (Kamei et al., 2011). Expression was verified by GFP fluorescence. Larvae heat shocked on 5 dpf were tested for behavior 3-6 hours after heat shock. Induced expression of *dnIGF1Ra-EGFP* from 24-72 hpf caused 10-20% of the embryos to show mild morphological abnormalities and these embryos were removed and not tested for behavior. Onset of *dnIGF1Ra-EGFP* expression at 3 dpf and after did not yield morphological defects.

### Immunohistochemistry and *in situ* hybridization

For whole mount immunostaining, larvae at 4-5 dpf were fixed in 4% paraformaldehyde (diluted to 4% w/v in phosphate buffered saline (PBS) from 16% w/v in 0.1M phosphate buffer, pH 7.4) for 1 hour at room temperature. Larvae were permeabilized in collagenase (0.1% w/v in PBS) for 3-4 hours and blocked for 1 hour at room temperature in incubation buffer (IB: 0.2% w/v bovine serum albumin, 2% v/v normal goat serum, 0.8% v/v Triton-X, 1% v/v DMSO, in PBS pH 7.4). Larvae were incubated in primary antibodies (see below) in IB overnight at 4°C. Larvae were incubated in fluorescently conjugated secondary antibodies in IB for 3-4 hours at room temperature and kept covered to block exposure to light thereafter. To stain cell nuclei, larvae were incubated in DAPI (0.1mg/mL in PBST, Sigma) for 1 hour after the secondary antibody was removed. After staining, larval heads were dissected and mounted in Vectashield (Vector Labs), while the tails were used to genotype for *pappaa*^*p170*^. Images were acquired with an Olympus Fluoview confocal laser scanning microscope (FV1000) using Fluoview software (FV10-ASW 4.2). To determine relative fluorescent intensity, raw integrated densities were measured from regions of interest and background in summation projections (10 focal planes spanning 4.6μm) using ImageJ software (Schindelin et al., 2012). Regions of interest were drawn around the retinal IPL, retinal OPL, and tectum. Relative fluorescence was reported as the ratio to DAPI staining in adjacent cell layers in the photoreceptor layer, retinal ganglion cell layer, and tectum.

Primary antibodies included: red/green double cone photoreceptors (anti-zpr-1, 1:100, mouse IgG1, Abcam); rod photoreceptors (anti-4c12, 1:100, mouse IgG, gift of Dr. Jim Fadool); ON bipolar cells (anti-PKCα, 1:100, rabbit IgG, Santa Cruz Biotechnology); amacrine cells (anti-5E11, 1:50, mouse IgG, gift of Dr. Jim Fadool); horizontal cells (anti-prox1, 1:100, mouse IgG, MilliporeSigma); bipolar and horizontal cells (anti-Lin7, 1:100, rabbit polyclonal serum, gift of Dr. Xiangyun Wei); Müller glia (anti-zrf-1, 1:500, mouse IgG1, ZIRC); excitatory postsynaptic density (anti-pan-MAGUK clone K28/86, 1:100, mouse IgG1, UC Davis/NIH NeuroMab Facility); inhibitory postsynaptic density (anti-gephyrin, 1:100, mouse IgG1, Synaptic Systems); phosphorylated IGF1R (anti-IGF1 receptor phospho Y1161, 1:100, rabbit IgG, Abcam); AMPAR (anti-GluR4, 1:100, rabbit, MilliporeSigma); synaptic vesicle protein (anti-SV2, 1:100, mouse IgG1, DSHB); retinal ganglion cell projections and photoreceptors using larvae from *Tg(isl2b-gfp);pappaa*^*p170/+*^ intercrosses (anti-GFP, 1:500, rabbit IgG, ThermoFisher Scientific). Secondary antibodies included: AlexaFluor488 and AlexaFluor594 conjugated secondary antibodies (goat anti-mouse IgG and IgG1, goat anti-rabbit IgG, 1:500, ThermoFisher Scientific).

Whole mount *in situ* hybridization was performed on larva at 36, 72, and 120 hpf as previously described (Halloran et al., 1999; Chalasani et al., 2007b) using digoxygenin-UTP labeled antisense riboprobes for *pappaa* (Wolman et al., 2015). Images were acquired with an Olympus Fluoview confocal laser scanning microscope (FV1000) using Fluoview software (FV10-ASW 4.2).

### Histology

For light microscopy, larvae at 5 and 10 dpf were processed as previously described (Sullivan-Brown et al., 2011). Larvae were fixed in 4% paraformaldehyde (diluted to 4% w/v in phosphate buffered saline (PBS) from 16% w/v in 0.1M phosphate buffer, pH 7.4) overnight at 4°C, dehydrated in a graded series of ethanol, and embedded in paraffin wax. Sections were cut at 3μm. Nuclei were stained with Harris hematoxylin and cytoplasm was stained with eosin Y. Images were acquired on a Nikon inverted light microscope (Eclipse TE300) using SPOT software (v5.1, Diagnostic Instruments, Sterling Heights, MI, USA). Plexiform layer width and cell counts for each sample were measured from the histological section in which the lens was at its largest diameter. Plexiform layer width was averaged from 6 measurements each taken 10μm apart. Retinal cell bodies were counted across the entire section and averaged per area of the different retinal nuclear layers.

For electron microscopy, larvae at 5 dpf were processed as previously described (Allwardt et al., 2001). Larvae were fixed (1% paraformaldehyde, 1.6% glutaraldehyde, 0.15 mM CaCl_2_, and 3% sucrose in 0.06 M phosphate buffer, pH7.4) for 15 minutes at 4°C and then rinsed and post-fixed in osmium tetroxide (2% in phosphate buffer) for 30 minutes at 4°C and 1.5 hours at room temperature. The tissue was then rinsed in phosphate buffer and in maleate buffer (0.05 M, pH 5.9) before being processed in a solution of 2% uranyl acetate in maleate buffer. The tissue was dehydrated in a graded series of ethanol, immersed into propylene oxide for 20 minutes, infiltrated with Araldite/Epon resin, and cured for 72 hours at 60°C. Approximately 80nm thick sections were mounted on slot grids and post-stained with lead citrate and saturated uranyl acetate. Images were acquired on a Philips CM120 scanning transmission electron microscope with a BioSprint 12 series digital camera using AMT Image Capture Software Engine V700 (Advanced Microscopy Techniques, Woburn, MA). Image coloring to indicate cell type was added with Adobe Illustrator CC 2018 (Adobe Systems, San Jose, CA, USA).

### Electroretinogram

Electroretinography was performed on larvae at 5-6 dpf. Experiments were performed at room temperature. Larvae were anesthetized in tricaine (0.002% in E3, Sigma) for 1 minute and then immobilized in low melting point agar (3% with 0.002% tricaine in E3, Sigma) with one eye exposed. Larvae were placed in the recording chamber under a stimulus light (X-cite Exacte; Lumen Dynamics, Mississauga, Ontario, Canada; light output adjusted to 90 μW/cm^2^ with neutral density filters). A glass electrode with a tip diameter between 20-40μm and filled with E3 medium was placed against the cornea. A reference electrode was inserted into the agar. Larvae were allowed to acclimate for at least 3 minutes before stimuli presentation. Larvae were presented with 20 light offset stimuli (dark flashes) lasting 1 second with a 10 second interstimulus interval. Reported traces are averages from the 20 stimuli and graphed with MATLAB software (R2017a, Mathworks, Natick, MA). The voltage signal was amplified (2K gain, MultiClamp-700A, Molecular Devices, Sunnyvale, CA), band-pass filtered (0.1 Hz - 1KHz), and digitized (40kHz; Digidata 1322A, Molecular Devices). Wild type and *pappaa*^*p170*^ mutant larvae were identified behaviorally by an acoustic startle habituation assay at 4 dpf as previously described (Wolman et al., 2011; Wolman et al., 2015).

### Statistics

Statistical analyses, including calculation of means, standard error, and ANOVA were performed using Microsoft Excel (Microsoft Corporation, Redmond, WA, USA) and GraphPad Prism software (GraphPad Software Incorporated, La Jolla, CA, USA).

## Results

### *pappaa* mutants show impaired initiation of behaviors to light offset

By 5 days post fertilization (dpf), larval zebrafish perform a repertoire of stereotyped motor behaviors in response to changes in visual field illumination (Neuhauss, 2003; Burgess and Granato, 2007b; Portugues and Engert, 2009; Burgess et al., 2010). For example, after light adaptation, exposure to the sudden absence of light, termed a ‘dark flash’, elicits an O-bend response in which larvae perform a large angle turn and reorient their body position by ~180 degrees (Burgess and Granato, 2007b). To determine whether Pappaa activity is required for larvae to perform this response, we exposed *pappaa*^*p170*^ larvae to 10 dark flashes and examined the initiation and performance of their O-bends. *pappaa*^*p170*^ larvae showed a marked reduction in O-bend initiation compared to wild type and *pappaa*^*p170U*^ larvae (Fig. 1C). When*pappaa*^*p170*^larvae initiated an O-bend (n= 25 O-bends), their mean latency to initiate the O-bend *(pappaa*^*p170*^: 244.4 msec +/− 10.85 SEM; *pappaa*^*+/+*^: 238.7 msec +/− 6.34 SEM, p=0.74, T-test), their O-bends’ mean body curvature *(pappaa*^*p170*^: 173.2 deg +/− 4.68 SEM; *pappaa*^*+/+*^: 169 deg +/− 1.45 SEM, p=0.33, T-test), and their mean distance moved as a result of O-bending *(pappaa*^*p170*^: 7.70 mm +/− 0.63 SEM; *pappaa*^*+/+*^: 7.94 mm +/− 0.22 SEM, p=0.69, T-test) were indistinguishable from O-bends performed by wild type larvae (n= 166 O-bends). These results and prior characterization of the *pappaa*^*p170*^ mutants (Wolman et al., 2015) suggest motor dysfunction does not underlie the *pappaa*^*p170*^ mutants’ reduced O-bend response initiation. The probability of initiating an O-bend response is influenced by whether a larva has adapted to the illumination conditions prior to light offset (Burgess and Granato, 2007b). We therefore evaluated whether the *pappaa*^*p170*^ larvae were light adapted by quantifying their initiation of swim and turn bouts in the minutes following a dark to light transition. As previously described (Burgess and Granato, 2007b) and similar to wild type larvae, *pappaa*^*p170*^ larvae exhibited an increase in motor activity (Fig. 2A-B), indicative of normal light adaptation. Given the *pappaa*^*p170*^ larvae’s impaired responsiveness to light offset, we next asked whether Pappaa activity is also required for motor response initiation to sudden light onset. To address this, larvae were adapted to a dimly illuminated arena and then exposed to a sudden increase in illumination (a “light flash”), which elicits a low angle turn response (Burgess and Granato, 2007b). Wild type and *pappaa*^*p170*^ larvae showed a similar response frequency to the series of light flashes (Fig. 1D). Together, these results indicate that Pappaa is required for larvae to respond to light offset, but not light onset.

A distinction between light and dark underlies contrast perception, which zebrafish larvae use for navigation (Burgess et al., 2010). Given the *pappaa*^*p170*^ larvae’s impaired response initiation to light offset, we hypothesized the mutants had defective contrast perception. To test this hypothesis we performed an optokinetic reflex (OKR) assay, which measures eye tracking of rotating stripes that vary in contrast (Brockerhoff et al., 1995; Emran et al., 2008). At bars ranging in contrast from 0.2 to 1, *pappaa*^*p170*^ larvae showed a reduced ability to track the moving bars compared to wild type larvae (Fig. 2C). To evaluate whether contrast-mediated navigation was also affected in the *pappaa*^*p170*^ larvae we performed a phototaxis assay. Larvae were first adapted to uniformly illuminated arena and then presented with a small target light on one side of the arena, which was now otherwise darkened (Burgess et al., 2010). Under these conditions wild type larvae stereotypically initiate a turn away from the darkened side of the dish and then initiate swim bouts toward the target light (Burgess et al., 2010). Indeed, wild type larvae positioned perpendicular to the luminance gradient initiated a turn away from the darker side of the gradient (Fig. 2D). When similarly positioned, *pappaa*^*p170*^ larvae showed no directional bias in their turns, indicating a failure to turn away from darkness (Fig. 2D). However, when the *pappaa*^*p170*^ larvae faced the target light, they initiated swim bouts toward the light similarly to wild type larvae (Fig. 2E). Together, these results suggest that Pappaa mediates the detection of light offset, which underlies contrast-dependent visually guided behavior.

### Pappaa acts through IGF1R signaling to promote behavioral responses to light offset

A role for Pappaa in detecting illumination change and mediating visually guided behavior is novel. Therefore, we sought to characterize the molecular pathway by which Pappaa promotes behavioral responses to light offset. Pappaa is a secreted metalloprotease that cleaves IGF binding proteins (IGFBPs) and therefore increases local IGF availability and activation of IGF receptors (Fig. 3A) (Lawrence et al., 1999; Conover, 2012; Oxvig, 2015). We first asked whether Pappaa’s canonical proteolytic activity towards IGFBPs is required for behavioral responses to light offset. To address this, we tested whether an isoform of human PAPPA lacking protease activity *(h-pappa^E483A^)* could improve the initiation of O-bend responses in *pappaa*^*p170*^ larvae (Boldt et al., 2001). Control injections of *h-pappa* mRNA into one-cell stage embryos significantly increased O-bend initiation by *pappaa*^*p170*^ larvae (Fig. 3B). However, injections of equimolar amounts of *h-pappa^E483A^* mRNA into*pappaa*^*p170*^did not improve their initiation of O-bends. These results indicate that Pappaa’s metalloprotease activity is required for initiating responses to light offset.

We next asked whether Pappaa acts via IGF1R signaling to promote light offset responses. We hypothesized that if Pappaa acts through IGF1R signaling, then attenuating IGF1R activity would reduce O-bend response initiation. To test this hypothesis, we used a *Tg(hsp70:dnIGF1Ra-GFP)* transgenic line, in which a heat shock induces ubiquitous expression of dominant negative *IGF1Ra* for approximately 12 hours (Kamei et al., 2011). *Tg(hsp70:dnIGF1Ra-GFP)* larvae heat shocked once every 12 hours from 1 to 5 dpf showed reduced initiation of O-bends (Fig. 3C). This reduction by *Tg(hsp70:dnIGF1Ra-GFP)* expression was stronger in the *pappaa*^*p170U*^ background (Fig. 3D), suggesting a genetic interaction between *pappaa* and *igf1ra.* As controls, nontransgenic larvae were similarly heat shocked and *Tg(hsp70:dnIGF1Ra-GFP)* larvae were not subjected to heat shock. Both groups exhibited normal O-bend responsiveness (Fig. 3C-D). Next, we hypothesized that if Pappaa acts through IGF1R signaling, then stimulating either IGF1 availability or a downstream effector of the IGF1R would improve O-bend responsiveness in *pappaa*^*p170*^ larvae. To test this hypothesis, we bathed wild type and *pappaa*^*p170*^ larvae in recombinant human IGF1 protein or a small molecule activator of Akt (SC79), a canonical downstream effector of IGF1R signaling (Laviola et al., 2007; Jo et al., 2012). These treatments from 1 to 5 dpf improved O-bend responsiveness in *pappaa*^*p170*^ larvae in a dose-dependent manner (Fig. 3E, G). Together, these results suggest that Pappaa acts through IGF1R signaling to promote a behavioral response to light offset.

We then addressed when Pappaa dependent IGF1R signaling was required to promote O-bend response initiation. First, we hypothesized that the minimal period of *Tg(hsp70:dnIGF1Ra-GFP)* expression required to reduce O-bend responsiveness would define the interval of IGF1R signaling that mediates O-bend initiation. Induction of *dnlGFlRa-GFP* expression that was restricted to 1-3 dpf or only at 5 dpf was insufficient to strongly reduce O-bend initiation (Fig. 3C-D). Rather, we found that *dnIGF1Ra-GFP* expression from 3-5 dpf reduced O-bend initiation to a degree that most closely resembled the O-bend reduction caused by continuous *dnIGF1Ra-GFP* expression from 1-5 dpf (Fig. 3C-D). Next, we hypothesized that the minimal period of IGF1 or SC79 exposure sufficient to improve O-bend responsiveness in *pappaa*^*p170*^ larvae would define the critical period of Pappaa-IGF1R-Akt signaling. *pappaa*^*p170*^ larvae exposed to IGF1 or SC79 only from 1-3 dpf or not until 5 dpf failed to improve their O-bend initiation (Fig. 3F, H). In contrast, IGF1 or SC79 treatments between 3-5 dpf significantly increased *pappaa*^*p170*^ larvae’s O-bend responses (Fig. 3F, H). Together, these results suggest that Pappaa-IGF1R-Akt signaling acts from 3-5 dpf to promote a larvae’s response to light offset.

### *pappaa* is expressed in the retinal inner nuclear layer

The behavioral analyses of the *pappaa*^*p170*^ larvae suggest that *pappaa* activity mediates the detection and interpretation of light offset, which occurs in the larvae’s retina and its projection targets. In mammals, retinal expression of PAPPA has been reported (Kay et al., 2012), but not described in temporal or spatial detail. We analyzed *pappaa mRNA* expression by whole-mount *in situ* hybridization from 36 to 120 hpf, in both the retina and its primary projection area: the optic tecta. This period encompasses when retinal and tectal neurons differentiate, form synapses, and establish the circuits that are sufficient to drive visually guided behaviors in 5 dpf larvae. We observed transient *pappaa* expression in the ventral retina at 36 hpf and in the temporal region of the retinal ganglion cell layer at 72 hpf (Fig. 4A-B). Beginning at 72 hpf and persisting through 120 hpf, *pappaa* was expressed in the retinal inner nuclear layer (INL) where the bipolar cells’ somas are positioned (Fig. 4B-C, bracketed). *pappaa* expression was not observed within the optic tecta between 72-120 hpf (Fig. 4D-E), although *pappaa* expression was detected in the brain ventral and medial to the tecta. Thus, *pappaa* is expressed in the distal retinal INL when synapses form between cones and bipolar cells. At this time and consistent with IGF1R signaling mediating Pappaa dependent responses to light offset (Fig. 3), we visualized activated IGF1R where cones and bipolar cells form synaptic connections in the outer plexiform layer (OPL) (Fig. 5A). Notably, we observed a reduction in anti-phosphorylated IGF1R (pIGF1R) immunolabeling in the OPL of *pappaa*^*p170*^ larvae (Fig. 5B-C). pIGF1R labeling of the inner plexiform layer (IPL) and in the tecta was similar in wild type and *pappaa*^*p170*^ larvae (Fig. 5C).

**Figure 4.**
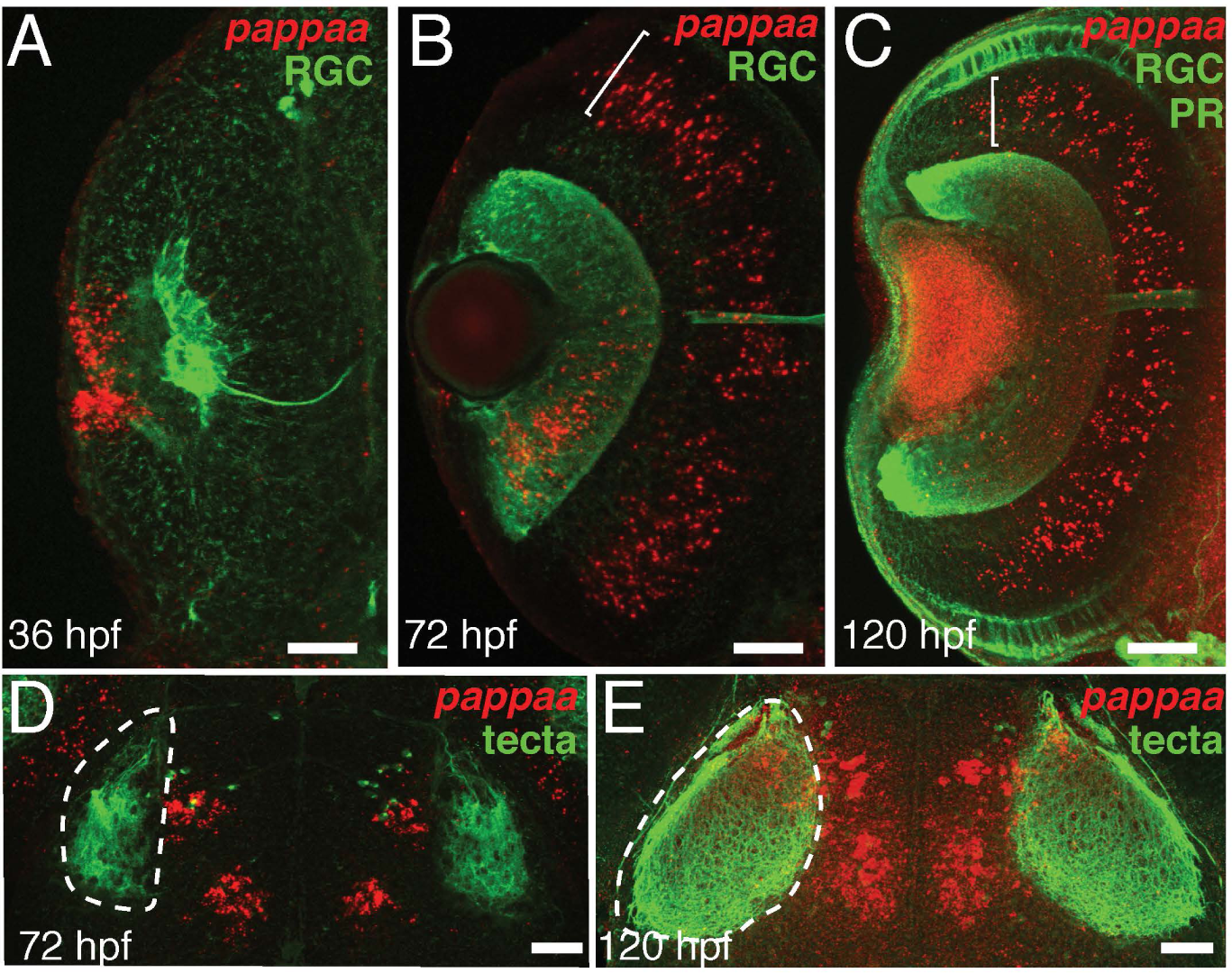
*pappaa* expression in the retina. *ln situ* hybridization for *pappaa* mRNA (red) in the retina at 36 (A), 72 (B), and 120 hpf (C) and in the tectal region (dashed line marks tectal boundary) at 72 (D) and 120 hpf (E). Bracket indicates retinal inner nuclear layer. All images are confocal projections of dorsal views, with anterior to the top of each panel. *Tg(lsl2b:GFP)* marks the retinal ganglion cells (RGC, A-C), photoreceptors (PR, C), and RGC axons’ innervation of the tecta (D-E). Labeling of lens in C is autofluorescence. Scale bar = 50 μm.

**Figure 5.**
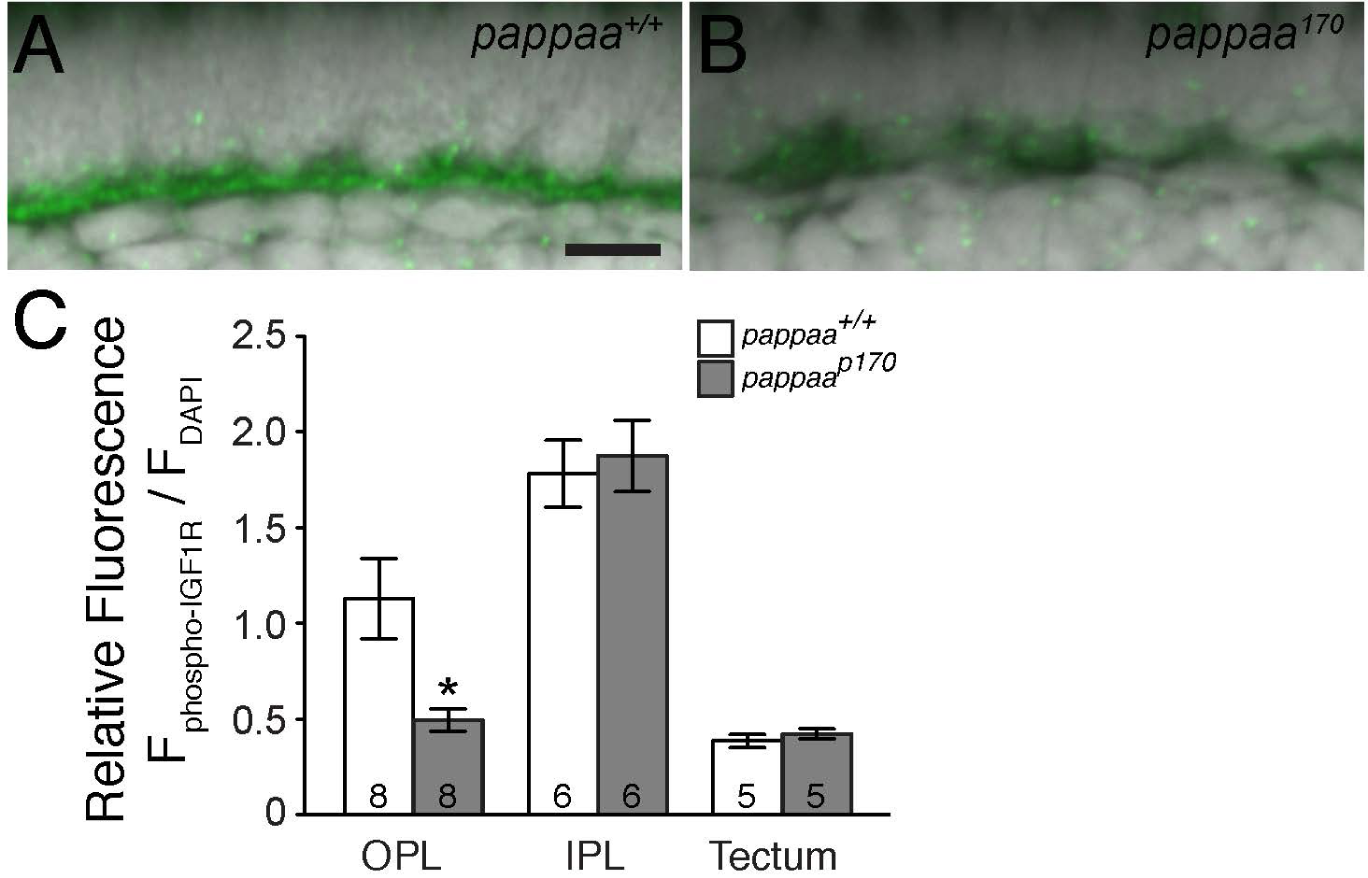
Phosphorylated IGF1R immunolabeling of the outer plexiform layer. (A-B) Summation intensity projections of anti-phosphorylated IGF1R immunolabeling of the OPL in wild type (A) and *pappaa*^*p170*^ larvae (B) at 5 dpf. Scale bar = 5 μm. (C) Mean fluorescent intensity of anti-phosphorylated IGF1R within OPL. (D) Ratio of mean fluorescent intensity of anti-phosphorylated IGF1R to mean fluorescent intensity of DAPI labeled photoreceptors for OPL, inner nuclear layer cells for INL, or tectal neurons for tecta. p<0.05 T-test vs wild type. N larvae shown at base of bars. Error bars indicate SEM.

### *pappaa*^*p170*^ retinas have a disrupted outer plexiform layer

*pappaa’s* earlier (36 hpf) retinal expression suggested that Pappaa loss of function could have widespread effects on retinal development that might disrupt intraretinal processing of light offset. To assess this possibility, we evaluated retinal lamination and gross cytoarchitecture by performing histology on wild type and *pappaa*^*p170*^ retinas at 5 and 10 dpf. Consistent with the smaller size of *pappaa*^*p170*^ larvae and growth promoting role of Pappaa-IGF1 signaling (Conover et al., 2004; Wolman et al., 2015), the *pappaa*^*p170*^ retinas appeared slightly smaller in size. By H&E staining, *pappaa*^*p170*^ retinas showed overtly normal lamination (Fig. 6A-B, E-F) and the number of cells per area in each retinal layer were similar between wild type and *pappaa*^*p170*^ larvae (Fig. 6J). To determine if each layer consisted of the proper retinal cell types, we performed whole mount immunolabeling with markers of cone and rod photoreceptors, bipolar cells, horizontal cells, amacrine cells, radial glia, retinal ganglion cells, and the retinotectal projection. All of these cell types were present in *pappaa*^*p170*^ retinas, in the proper layer, and with the same abundance as observed in wild type retinas (Fig. 7A-H’). The retinotectal projections appeared normal in *pappaa*^*p170*^ larvae, as indicated by anatomy of the projection (Fig. 7H-H’) and the degree of tectal innervation at 5 dpf (Fig. 7I-I’; *isl2b-gfp* fluorescent intensity, raw integrated density per area (units/μm^2^) minus background: wild type = 78,250.97 +/− 6,228.36 SEM, n = 10 samples, 1 tectum per sample; *pappaa*^*p170*^ = 93,531.96 +/− 4,294.35 SEM, n = 11 samples, 1 tectum per sample; p = 0.06, T-test). A closer inspection of the 5 dpf retinal histology revealed that the width of the *pappaa*^*p170*^ larvae’s OPL was reduced by approximately 12.66% (Fig. 6C-D, I). At 10 dpf, the *pappaa*^*p170*^ larvae’s OPL appeared disorganized and the reduction in width was more pronounced (38.81%, Fig. 6G-I), indicating the difference at 5 dpf was not transient and that the phenotype progressively worsened. Notably, the width of the IPL was equivalent between genotypes at both stages (Fig. 6I).

**Figure 6.**
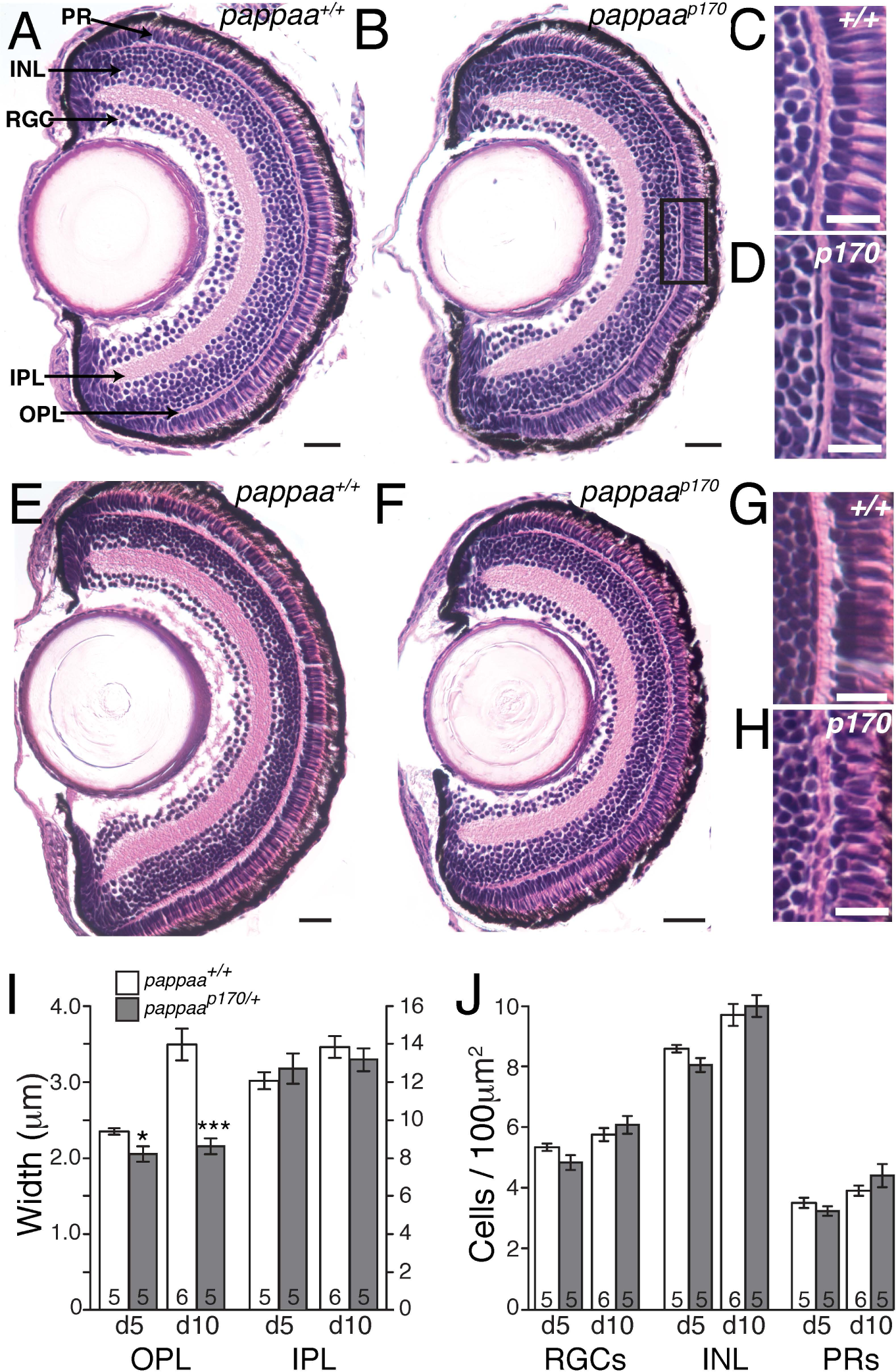
Cytoarchitecture of *pappaa*^*p170*^ retina. Histological sections of retinas at 5 (A-D) and 10 dpf (E-H) retinas of wild type and *pappaa*^*p170*^ larvae, stained with H&E. Scale bars = 20μm in A-B, E-F; and 10μm in C-D, G-H. (I) Mean width of OPL and IPL. (J) Mean counts of cells in retinal ganglion cell layer (RGC), inner nuclear layer (INL), and photoreceptor layer (PR). N = number of larvae shown at base of bars. *p<0.05, ***p<0.001 vs. wild type at same stage, Students T-test. Error bars indicate SEM.

**Figure 7.**
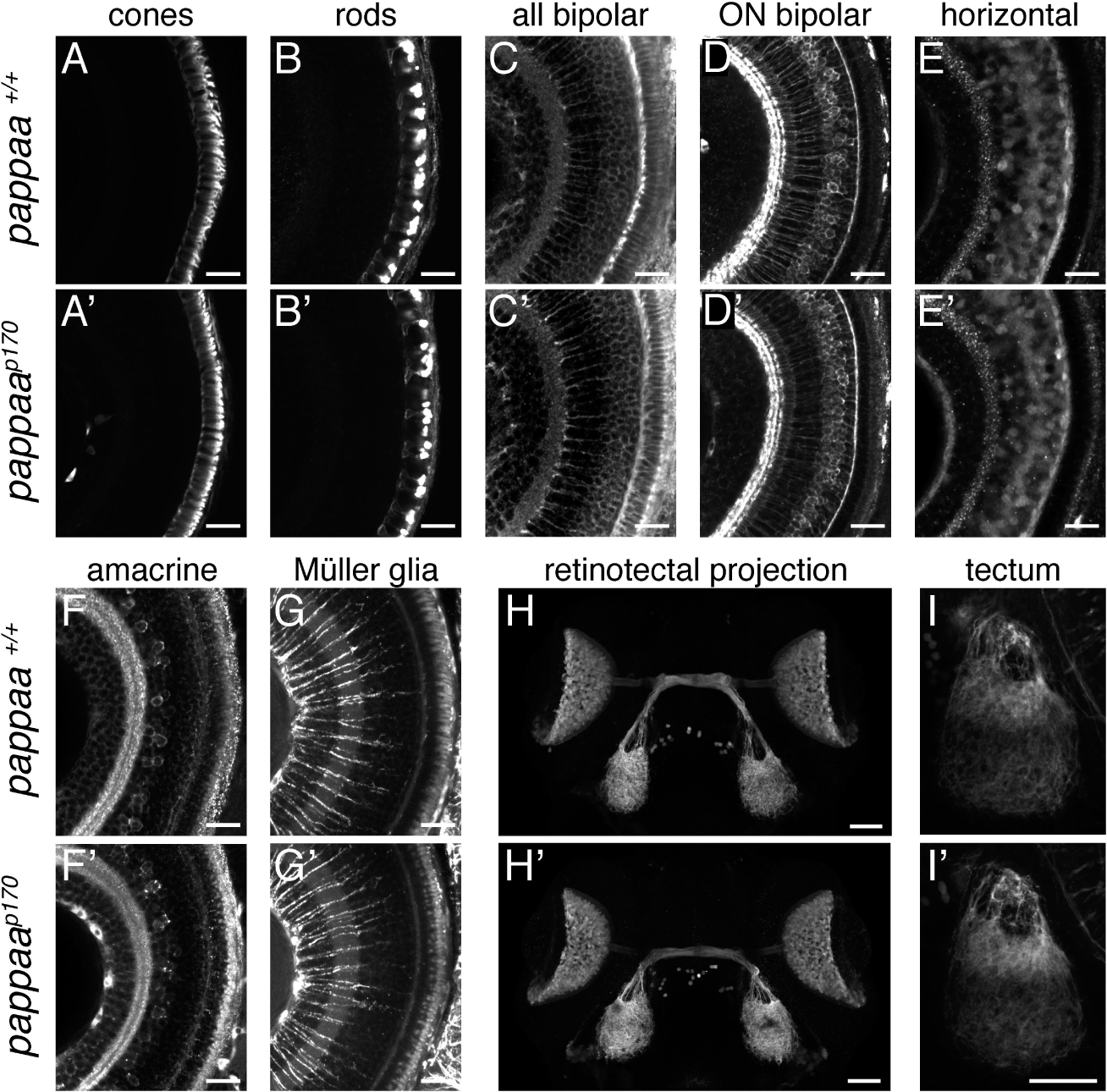
Retinal cell types and retinotectal projection of *pappaa*^*p170*^ larvae. All images are maximal confocal projections (unless otherwise noted) of dorsal views, anterior to the top. (A-G’) Whole mount immunolabeling of wild type and*pappaa*^*p170*^ larvae retinas for cone photoreceptors (zpr-1), rod photoreceptors (4c12), bipolar cells (Lin7), ON bipolar cells (PKCa), horizontal cells (proxl), amacrine cells (5E11), and Müller glia (zrf-1) at 96 hpf. (H-H’) Retinal ganglion cell (RGC) projections to tecta of 96 hpf wild type and *pappaa*^*p170*^ larvae as marked by *Tg(lsl2b:GFP).* (I-I’) Representative summation projections of RGC axonal innervation of tecta at 96 hpf in wild type and *pappaa*^*p170*^ larvae. RGC axons marked by *Tg(lsl2b:GFP).* Scale bars = 20μm (A-G’) and 50μm (H-I’).

A thin OPL has been associated with impaired photoreceptor synaptic development and visual function (Jia et al., 2014), and thus prompted an evaluation of synaptic markers in wild type and *pappaa*^*p170*^ retinas. We labeled 5 dpf retinas with antibodies against postsynaptic scaffolding proteins that mark excitatory (anti-MAGUK) and inhibitory (anti-gephyrin) postsynaptic densities within both plexiform layers. In both genotypes, the OPL and IPL showed a similar degree of immunolabeling and the anti-gephyrin labeling marked clearly defined laminae within the IPL (Fig. 8A-D). Given the *pappaa*^*p170*^ larvae’s impaired responses to light offset, we next evaluated the expression of AMPA-type glutamate receptors on OFF-bipolar cells’ dendrites, which mediate their activation by glutamate release from photoreceptors at light offset. The number of anti-AMPA-R labeled puncta per cone in the OPL was equivalent between wild type and *pappaa*^*p170*^ larval retinas (Fig. 8E-F, wild type = 2.02 +/− 0.13 SEM, N = 8 samples, at least 15 cones per sample; *pappaa*^*p170*^ = 2.02 +/− 0.07 SEM, N = 8 samples, at least 14 cones per sample; p = 0.99, T-test). We next analyzed presynaptic vesicles in photoreceptors at the OPL by immunolabeling with anti-SV2. Although there was labeling in the OPL of both wild type and *pappaa*^*p170*^ larvae (Fig. 8G-H), we observed abundant irregularities in the anti-SV2 labeling in 75% of *pappaa*^*p170*^ larvae (N = 8), which were rarely observed in the OPL of wild type retinas. These irregularities included gaps in the anti-SV2 labeling (Fig 8H, arrowhead) and labeling that extended further into the photoreceptor layer than observed in wild type (Fig. 8H, arrows).

**Figure 8.**
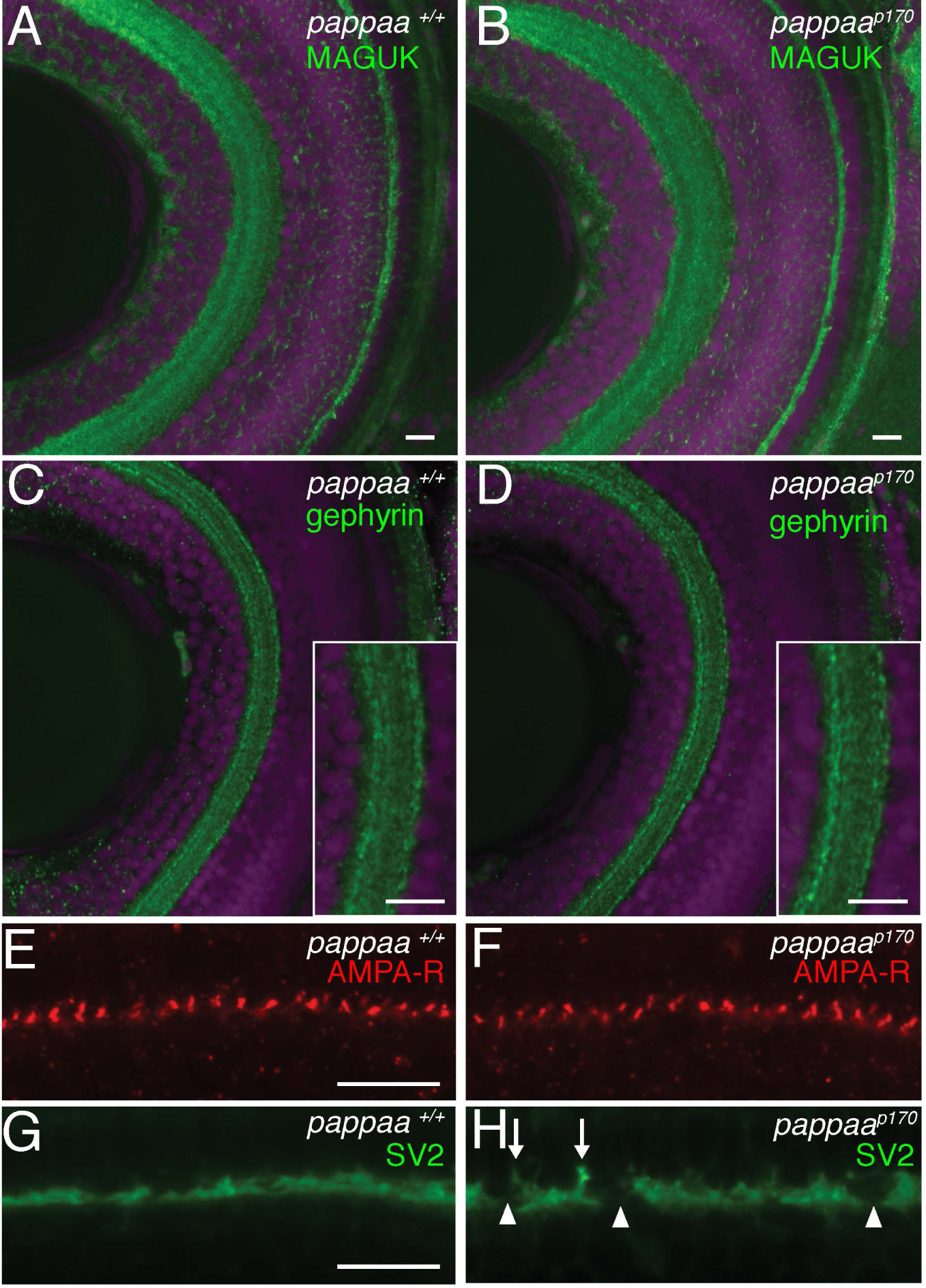
Retinal synaptic markers. Dorsal views, anterior to the top of 5 dpf wild type and *pappaa*^*p170*^ retinas labeled with anti-MAGUK (A-B) or anti-gephyrin antibodies (C-D) to mark excitatory and inhibitory post-synaptic densities, respectively. Insets in C-D show lamination of IPL. (E-F) AMPA receptors on OFF-bipolar cell dendrites labeled by anti-GluR4 antibody. (G-H) Synaptic vesicles in photoreceptors labeled by anti-SV2 antibody. Arrowheads mark gaps in anti-SV2 labeling rarely observed in wild type. Arrows mark anti-SV2 labeling that extended more proximally within the photoreceptor layer than observed in wild type. All scale bars = 10μm.

### *pappaa* mutants exhibit reduced cone to OFF bipolar cell synaptic activity

Cone photoreceptors modulate synaptic vesicle release within the OPL to activate parallel circuits that convey light onset versus offset (Fig. 1A-B). To light onset, cones hyperpolarize and reduce synaptic vesicle release to activate ON bipolar cells. To light offset, cones depolarize and increase synaptic vesicle release to activate OFF bipolar cells. Based on the *pappaa*^*p170*^ larvae’s behavioral defects being limited to light offset, we hypothesized that Pappaa activity is required for synaptic transmission between cones and OFF bipolar cells, but not ON bipolar cells. To measure outer retina synaptic function in response to both light offset and onset, we performed electroretinograms (ERGs) in immobilized larvae that were light adapted and then exposed to a series of 1s dark flashes. ERGs record neuronal field potential changes specific to ON or OFF bipolar cells following changes in luminance (Wong et al., 2004). Upon light offset 100% of ERG recordings from wild type larvae (n = 6 larvae, 20 stimuli per larva) showed an expected increase in amplitude (d-wave) (Fig. 9), which indicated functional synapses between cones and OFF bipolar cells. In contrast, ERG recordings from only 58% of *pappaa*^*p170*^ larvae (n = 12 larvae, 20 stimuli per larva) showed a typical d-wave. At light offset, the ERGs from 3/12 larvae had minimal amplitude change (Fig. 9B, single asterisk) and 2/12 larvae exhibited an inverted waveform (Fig. 9B, double asterisk). To the light onset that followed each 1s dark flash, ERG recordings from 100% of the wild type and *pappaa*^*p170*^ larvae exhibited the expected robust increase in amplitude (b-wave) (Fig. 9), which is indicative of functional cone phototransduction and synaptic transmission between cones and ON bipolar cells. These results suggest that *pappaa*^*p170*^ larvae have functionally normal cone to ON bipolar cell synapses, but have deficient synaptic activity between cones and OFF bipolar cells.

**Figure 9.**
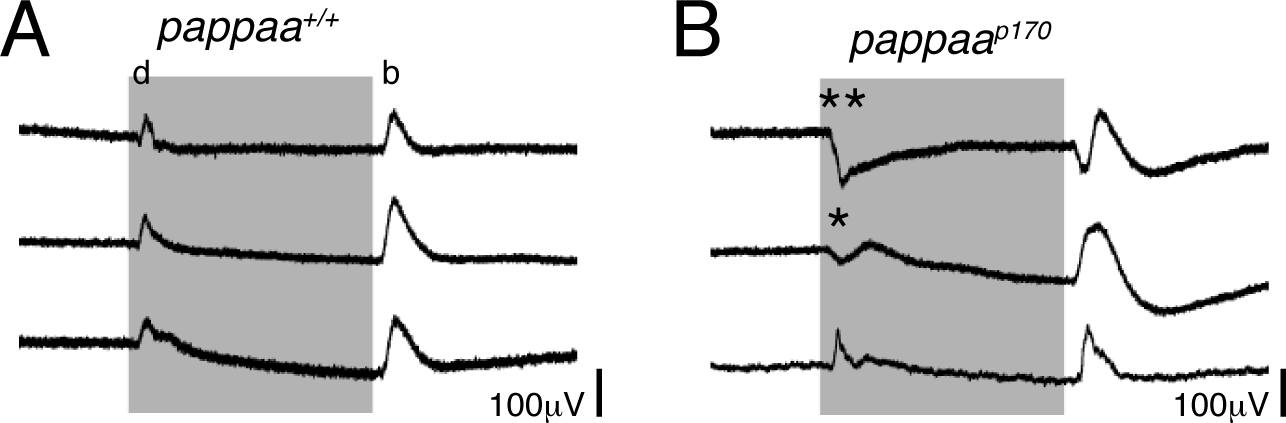
Electroretinograms wild type and *pappaa*^*p170*^ larvae. Electroretinogram recordings from 5 dpf wild type (A) and *pappaa*^*p170*^ larvae (B) to 1 sec light offset (grey region) and onset. Each trace is from a single larva and represents the mean change in amplitude from 20 recordings. All wild type traces showed increased amplitudes at light offset (d-wave) and onset (b-wave). For *pappaa*^*p170*^ larvae, three representative traces are shown for the three outcomes observed for the d-wave: * minimal d-wave, ** = negative d-wave, and a wild type-like d-wave (bottom).

### Pappaa-IGF1 signaling regulates presynaptic architecture of cone photoreceptors

We next evaluated the ultrastructural integrity of the OPL by performing electron microscopy (EM) on 5 dpf wild type and *pappaa*^*p170*^ retinas. Invaginations into the photoreceptor layer by bipolar and horizontal cell dendrites clearly marked regions within the OPL where cone photoreceptors formed synapses with these second order neurons (Fig. 10A). By comparing EM micrographs of wild type to *pappaa*^*p170*^, we observed a similar number of invaginations per 100μm along the anterior-posterior axis of the OPL (wild type = 15.47 +/− 0.84 SEM, n = 5 larvae; *pappaa*^*p170*^ = 14.53 +/− 0.94 SEM, n = 5 larvae; p = 0.53, T-test). These invaginations were of the same size (invagination area (μm^2^): wild type = 1.34 +/− 0.06 SEM, n = 5 larvae averaged from 10 invaginations per larva; *pappaa*^*p170*^ = 1.24 +/− 0.09 SEM, n = 5 larvae averaged from 10 invaginations per larva; p = 0.44, T-test). And, the invaginations consisted of horizontal and bipolar cell dendrites with equal abundance (Fig. 10B,D). Wild type and *pappaa*^*p170*^ cones possessed approximately the same number of synaptic vesicles within 150nm of the cone presynaptic membrane (vesicles per area (μm^2^): wild type = 126.40 +/− 4.02 SEM, n = 4 larvae, averaged from 3 invaginations per larva; *pappaa*^*p170*^ = 122.78 +/− 5.34 SEM, n = 4 larvae averaged from 3 invaginations per larva; p = 0.82, T-test). Next, we evaluated the cone presynaptic domains that are positionally coupled to ON and OFF bipolar cell dendrites. Flat contacts between cones and OFF bipolar cell dendrites were identified based on the apposed, electron-rich pre-and postsynaptic densities within the peripheral regions of the invaginations (Fig. 10A-C) (Dowling and Boycott, 1966; Nelson and Connaughton, 1995). In *pappaa*^*p170*^ larvae, we observed a striking 47.62% reduction in the number of invaginations with an identifiable flat contact (Fig. 10D-F). At contacts with ON bipolar cells, synaptic ribbons were highly visible, docked to the presynaptic cone membrane, and anchored by an archiform density. In *pappaa*^*p170*^ cones we observed an increase in the presence of “floating”, undocked ribbons that lacked an archiform density (Fig. 10G, asterisk); and thus, the percentage of docked ribbons was mildly reduced in *pappaa*^*p170*^ (Fig. 10G). To determine if Pappaa acts through IGF1 signaling to promote the integrity of these structures, we asked whether stimulation of IGF1 signaling in *pappaa*^*p170*^ larvae would restore these structures as it did for light offset responses. Indeed, exposure of *pappaa*^*p170*^ larvae to IGF1 from 3-5 dpf increased the percentage of invaginations with an identifiable flat contact and docked synaptic ribbons to wild type levels (Fig. 10F-G).

**Figure 10.**
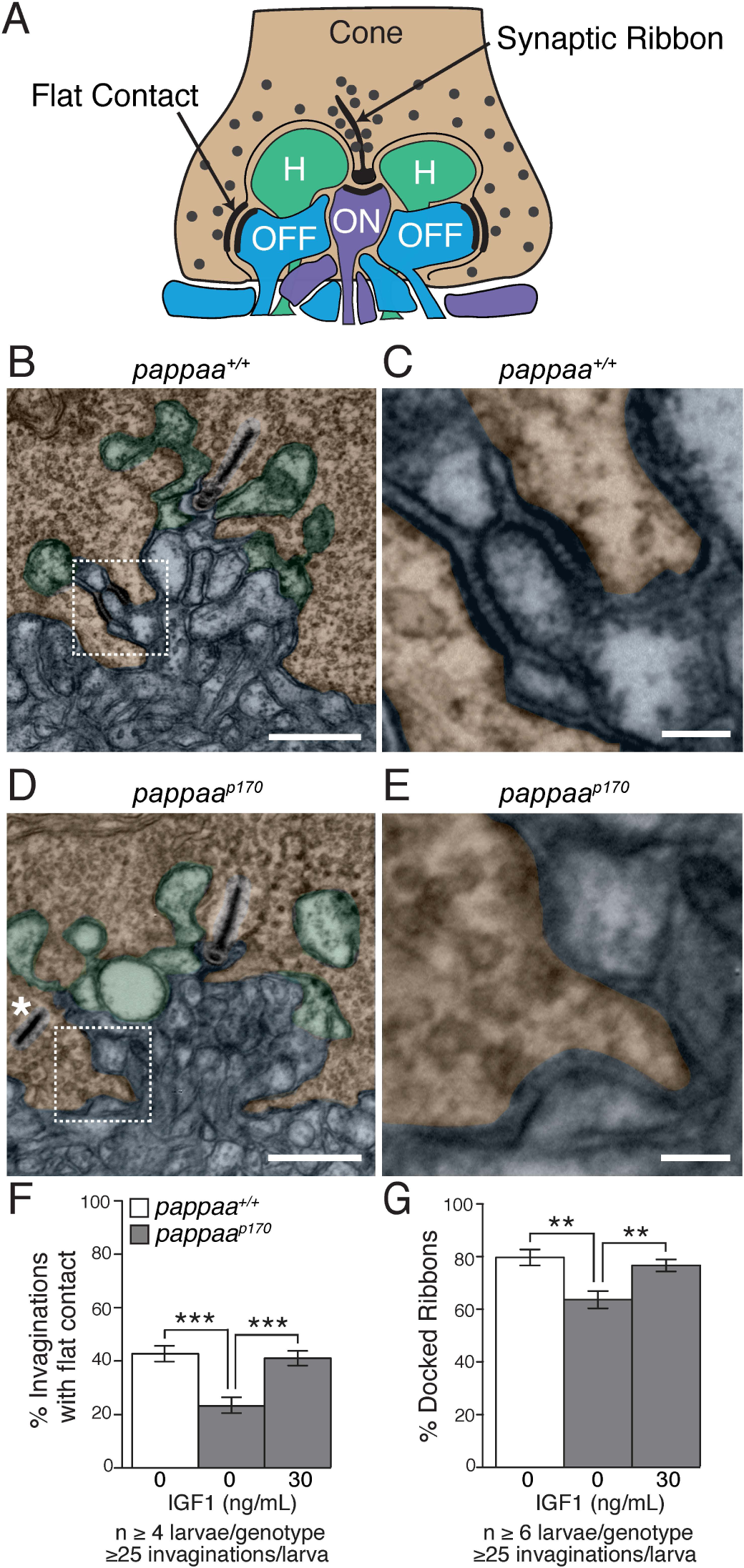
*pappaa*^*p170*^ larvae show reduced synaptic vesicle docking in retinal photoreceptors. (A) Schematic of photoreceptor synaptic terminal with dendrites of ON (purple) and OFF (blue) bipolar cells and horizontal (H, green) cells forming invaginations into the cone synaptic terminal. (BE) Electron micrographs of cone terminals in wild type (B, C) and *pappaa*^*p170*^ (D-E). Coloring matches cartoon in A, except all bipolar cell dendrites are blue-grey. (C, E) Higher magnification of boxed regions in B and D to highlight flat contact. Asterisk in D marks undocked, “floating” synaptic ribbons. Scale bars = 500nm (B, D) and 100nm (C, E). (F) Mean percentage of invaginations with an identifiable flat contact, as indicated by a pre-synaptic electron dense region found laterally within an invagination and across the synaptic cleft from a post-synaptic electron dense band. (G) Mean percentage of ribbons that were docked, as defined by proximity to cone’s presynaptic membrane and presence of archiform density. **p<0.01, ***p<0.001, One-way ANOVA. N indicated under graphs. Error bars indicate SEM.

The physiological role and structural makeup of flat contacts remains unclear. These domains have been dismissed as sites of synaptic vesicle release based on failure to visualize nearby and/or docked synaptic vesicles. Notably, our EM images indicated synaptic vesicles near the flat contacts’ presynaptic density (Fig. 11, arrowheads) (mean synaptic vesicles within 200nm = 4.92 +/− 1.20 SEM, N= 12 flat contacts from 4 wild type retinas). Consistent with an idea that loss of flat contacts’ presynaptic domain, and hence, reduced vesicle release may underlie the *pappaa*^*p170*^ larvae’s defect in responding to light offset, we found that acute, 20 minute treatment of the *pappaa*^*p170*^ larvae with AMPA was sufficient to restore their initiation of light offset responses (Fig. 12). Finally, we evaluated OFF bipolar cells’ presynaptic terminals in the IPL, since their loss could contribute to the *pappaa*^*p170*^ larvae’s impaired responses to light offset. Quantification of synaptic ribbons in the IPL, and in particular the outer laminae of the IPL where OFF bipolar cells project (Nelson and Connaughton, 1995; Nevin et al., 2008), did not reveal a difference between wild type and *pappaa*^*p170*^ (Fig. 13). Together, these results reveal a novel role for Pappaa regulated IGF1R signaling in supporting cone presynaptic structures.

**Figure 11.**
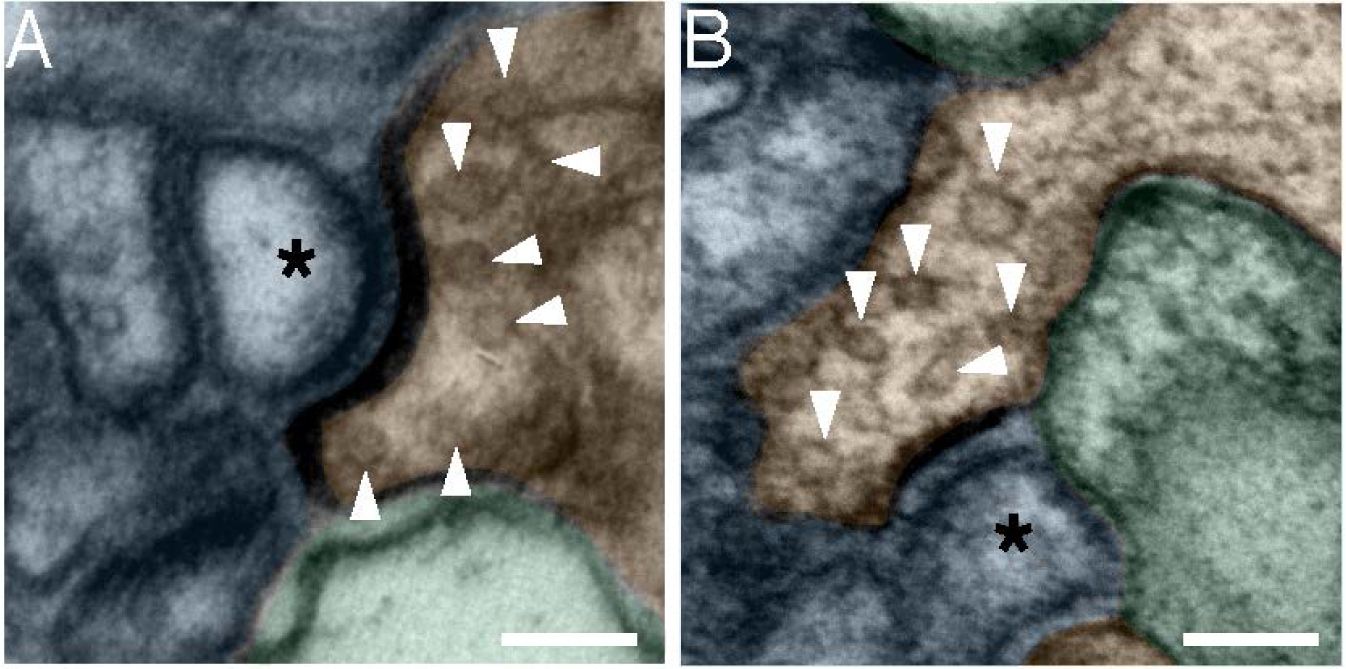
Synaptic vesicles are present at flat contacts. Electron micrographs showing examples of flat contacts from two wild type retinas. Arrowheads mark synaptic vesicles in cone (shaded brown). Bipolar dendrites are shaded blue grey. OFF bipolar dendrites are identified by a postsynaptic density of flat contact and are marked with asterisk. Horizontal cell dendrites shaded green. Scale bars = 100nm.

**Figure 12.**
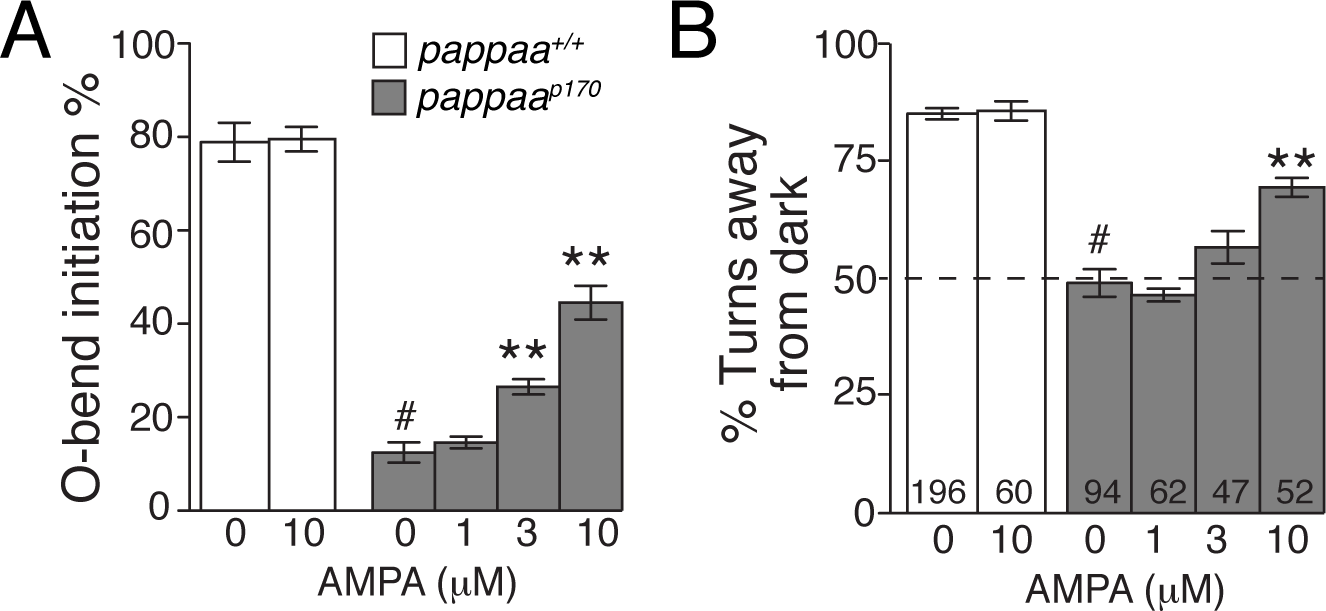
AMPA supplementation improves O-bend responsiveness in *pappaa*^*p170*^ larvae. Larvae were treated for 30 minutes prior to and during exposure to 10 dark flashes (A) or phototaxis assay (B) at 5 dpf. (A) Mean initiation frequency of O-bend responses. N = 5 groups of 15 larvae per treatment group. (B) Mean percentage of turns initiated away from darkness by larvae positioned between 75-105 degrees to the light-dark gradient. N at base of bars = number of trials in which a turn was evaluated based on larval position with respect to target light. #p<0.001 vs. untreated wild type larvae and **p<0.01 vs. untreated *pappaa*^*p170*^ larvae, ANOVA with Bonferroni correction.

**Figure 13.**
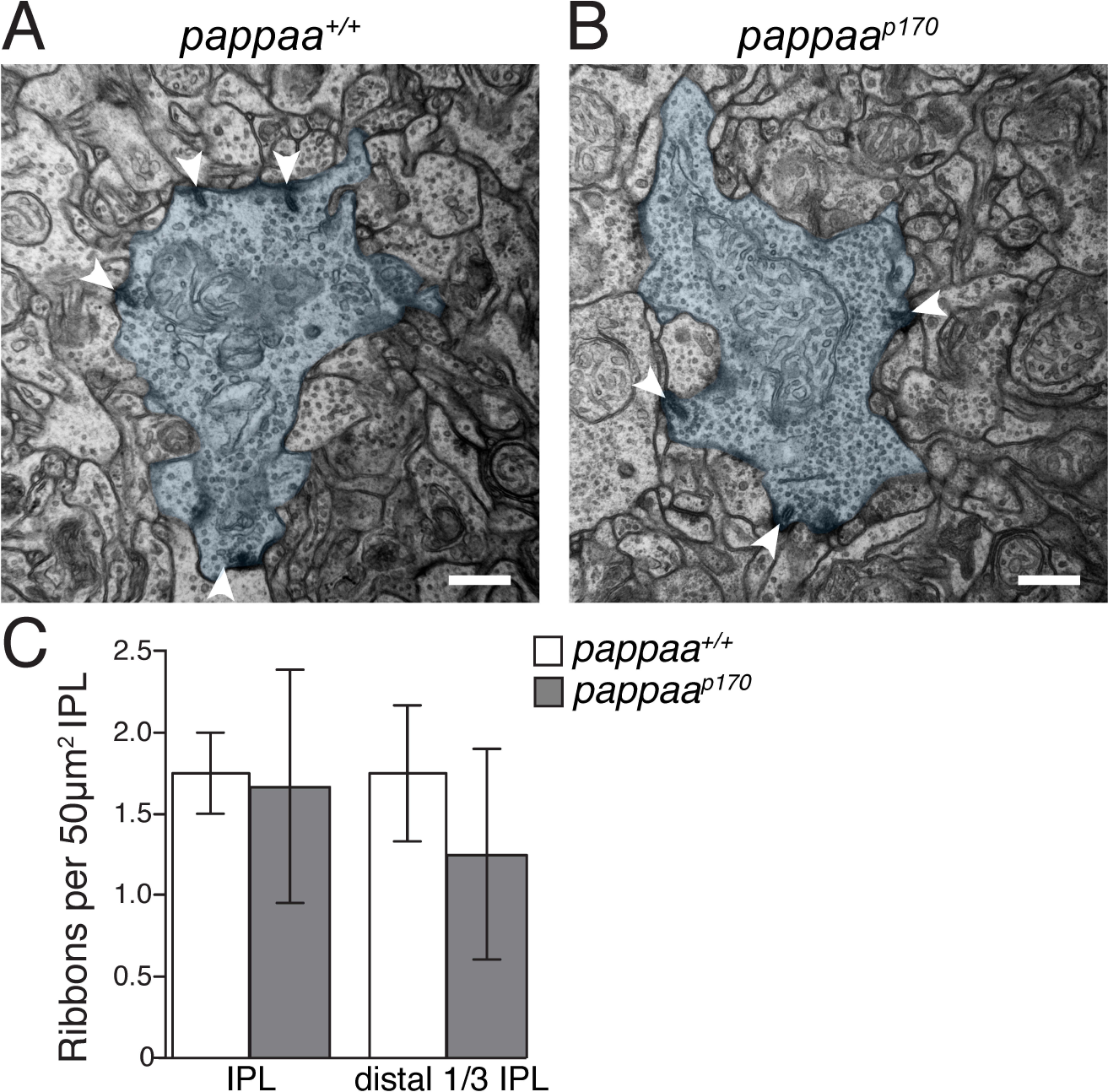
OFF bipolar cell synaptic ribbons in the IPL. Electron micrographs of IPL synapses in wild type (A) and *pappaa*^*p170*^ (B). OFF bipolar cell terminals (shaded in blue-grey) identified by location in the distal third of the IPL and presence of synaptic ribbons (white arrowheads). Scale bars = 500nm. (C) Mean number of synaptic ribbons per area of IPL. n = 4 larvae per genotype. Error bars indicate SEM.

## Discussion

Neural signals coding light onset and offset are split into parallel retinal circuits via discrete synapses between photoreceptors and distinct bipolar cells. Despite an understanding of these synapses’ physiology, it remains poorly understood how molecular cues establish their architecture to mediate signal splitting, particularly to convey light offset. Here, we characterized a unique zebrafish mutant with visual impairment specific to light offset. This analysis revealed Pappaa as novel molecular regulator of cone to OFF bipolar cell synaptic structure and function and describes Pappaa’s first known role in synapse development.

## Pappaa regulates visual responses specific to light offset stimuli

The precise impact of Pappaa on retinal circuits is demonstrated by the specificity of the *pappaa*^*p170*^ mutants’ defects in visually guided behaviors and cone to bipolar cell synaptic structure and physiology. *pappaa*^*p170*^ larvae failed to properly initiate behaviors to light offset (Fig. 1C, 2D), yet showed normal initiation of behaviors to light onset (Fig. 1D), during light adaptation (Fig. 2A-B), and when facing a light stimulus (Fig. 2E). When *pappaa*^*p170*^ larvae initiated behavior to light onset or offset, startled to sound, or moved spontaneously (Wolman et al., 2015), the kinematic parameters of their maneuvers were normal, indicating intact motor function. This analysis suggests Pappaa acts upstream of the visual circuit’s motor component to regulate detection and/or interpretation of light offset. The significance of this role was further demonstrated by the *pappaa*^*p170*^ larvae’s diminished contrast-dependent behaviors, including the OKR (Fig. 2C), phototaxis navigation (Fig. 2D-E), and prey capture (Miller and Wolman, unpublished).

Multiple observations suggest Pappaa acts in the outer retina to mediate light offset detection and/or interpretation. Processing of light input occurs in the retina and the retina’s projection targets in the brain, which is primarily the tectum in zebrafish. Within the timeframe of our analyses, *pappaa* was not expressed by the tecta, but was retinally expressed (Fig. 4). When IGF1 signaling was required for light offset responses (Fig. 3C-H), *pappaa’s* retinal expression was largely restricted to the INL (Fig. 4B-C), and thus, postsynaptic for cone to bipolar cell synapses. From ERGs recorded in *pappaa*^*p170*^ larvae, normal cone to ON bipolar cell waveforms indicated these synapses were functional and cone phototransduction was intact (Fig. 9). The aberrant waveforms at light offset revealed dysfunction specific to the cone to OFF bipolar cell synapses (Fig. 9B). Notably, the ERG revealed a milder defect in *pappaa*^*p170*^ larvae compared to their more robust behavioral deficit. This discrepancy could be due to the ERG being a less-sensitive assay for assessing light offset responses and/or to differences in how the larvae are tested by ERG and for behavior (ERG: immobilized, anesthetized versus behavior: free swimming) and/or the light stimuli used. Additionally, *pappaa*^*p170*^ larvae may harbor circuit defects downstream of the cone to OFF bipolar cell synapses, which would be undetectable by ERG, but could further disrupt transmission of the light offset neural signals. We failed to identify such a defect through our histological, ultrastructural, and immunohistochemical analyses. Rather, the structural defects we identified were limited to the cone presynaptic terminals.

## Pappaa regulates cone presynaptic structure

To understand how Pappaa promotes light offset responses it is important to consider where Pappaa influences retinal circuitry. *pappaa’s* INL expression (Fig. 4B-C) combined with the *pappaa*^*p170*^ larvae’s ERG (Fig. 9B), retinal histology (Fig. 6), and IGF1R phosphorylation (Fig. 5), together suggest Pappaa influences circuits within the OPL. Here, the synaptic configuration between the photoreceptors and bipolar cells splits inputs of light onset and offset. At light offset, transient glutamate release from cones activates AMPA receptors on OFF bipolar cell dendrites to transmit a message of light offset to ganglion cells (Wassle, 2004). Thus, impairment in initiating light offset responses could be caused by a structural and/or functional defect at either the cone to OFF bipolar synapses or the OFF bipolar cells’ innervation of the IPL. Multiple lines of evidence suggest Pappaa does not mediate OFF bipolar cell structure or function within the timeframe of our analyses. First, the number and size of dendritic invaginations into cone terminals were normal in *pappaa*^*p170*^. Second, AMPA receptors, expressed by horizontal and OFF bipolar cell dendrites, showed normal expression in *pappaa*^*p170*^. Third, *pappaa*^p170^mutant retinas showed normal innervation of the IPL’s outer laminae where OFF bipolar cells form synapses with ganglion cells (Fig. 7C-C’) and normal synaptic ribbons (Fig. 13). Finally, *pappaa*^*p170*^ larvae showed improved responses to light offset stimuli when acutely treated with AMPA (Fig. 12), suggesting their OFF bipolar cells were competent. Together, these results suggest the circuit defect underlying the *pappaa*^*p170*^ larvae’s deficient light offset response is unlikely to be localized to OFF bipolar cells.

The *pappaa*^*p170*^ cones’ reduced flat contact presynaptic domains and mislocalized synaptic ribbons provides evidence that the circuit defect underlying the impaired light offset responses is in the cones’ presynaptic terminal. Although ribbons mediate glutamate release at light offset, mutant analyses have demonstrated the cones’ ribbons are dispensable for light offset responses in larval zebrafish (Allwardt et al., 2001; Emran et al., 2008). Photoreceptors have non-ribbon mediated sites of glutamate release (Chen et al., 2013), but the presynaptic domains of flat contacts have been dismissed as a release site based on descriptions, including in zebrafish, that claim this region is devoid of synaptic vesicles (Dowling and Boycott, 1966; Haverkamp et al., 2000; Allwardt et al., 2001; DeVries et al., 2006). Our electron micrographs rebut this claim by showing vesicle-like structures near and apparently docked at the flat contacts’ presynaptic membrane (Fig. 11). Though fewer synaptic vesicles were observed at the flat contacts’ presynaptic density compared to at ribbons, it has been shown that a single vesicle’s worth of glutamate is sufficient to depolarize an OFF bipolar cell (DeVries et al., 2006). Glutamate release at the flat contact is positionally more favorable to activate AMPA receptors on OFF bipolar cell dendrites compared to the ribbon-mediated glutamate release that occurs hundreds of nanometers away. The possibility that the *pappaa*^*p170*^ mutant has uncovered a novel role for the flat contacts in mediating vesicular glutamate release is exciting, but will require direct evidence of synaptic vesicle release at this locus to be substantiated. To understand Pappaa’s relationship with flat contacts and this locus’s potential role in glutamate release and light offset responses, it will also be critical to characterize the structural composition of the flat contacts presynaptic domain, which remains a mystery (Boycott and Hopkins, 1993; Tsukamoto and Omi, 2015).

## Pappaa regulates IGF1R signaling to promote light offset responses and synaptic structure

Presynaptic domains develop through the signaling of molecular cues, which may be expressed and act within the presynaptic neuron, postsynaptic neuron, or surrounding cells (Cohen-Cory, 2002; Shen and Cowan, 2010). IGF1 regulates circuit development and function by binding the IGF1R, which activates intracellular signaling cascades including the PI3K-Akt-mTOR pathway (Dyer et al., 2016; Nieto-Estevez et al., 2016). For cone to bipolar cell synapses, our data suggest *pappaa* is expressed postsynaptically (Fig. 4B-C), and stimulates local IGF1R signaling during cone synapse development to establish flat contacts and mediate light offset responses (Fig. 3, 5, 10F). IGF1 signaling promotes synapse formation and function through pre and postsynaptic mechanisms and given the IGF1R’s widespread neural expression during development (Torres-Aleman, 1999; Dyer et al., 2016; Nieto-Estevez et al., 2016), locally acting regulators of IGF1 signaling, like Pappaa, must be important for circuits.

Evidence presented here suggests Pappaa-IGF1 signaling influences cone presynaptic domains at synapses with OFF bipolar cells. However, it remains unclear which outer retinal cell types require IGF1R activity to establish these domains. Pappaa could act in an autocrine manner to stimulate postsynaptic IGF1R signaling in OFF bipolar cells. Postsynaptically, IGF1 has been shown to promote dendritic growth (Camarero et al., 2001; Cheng et al., 2003; Shi et al., 2005; Carlson et al., 2014), membrane receptor and channel stability through the activity of synaptic scaffolding proteins (Corvin et al., 2012; Della Sala et al., 2016), and the development and maintenance of synaptic connections via cell adhesion molecules (Bonfanti, 2006; Monzo et al., 2013). Although we did not observe defects in dendritic size (Fig. 10B, D) or abundance of AMPA receptor or postsynaptic density markers in the OPL of *pappaa*^*p170*^ mutant (Fig. 8), IGF1 signaling may act through other postsynaptic mechanisms to promote the development or maintenance of cone-OFF bipolar cell synaptic contacts. Alternatively, Pappaa may act in a paracrine manner to enhance IGF1R signaling in cones or another local cell type (e.g. horizontal cells or the still quiescent rods). Pappaa may free IGF1 to stimulate presynaptic, cone photoreceptor-dependent cellular mechanisms. IGF1 signaling has been shown to regulate axon guidance (Laurino et al., 2005; Scolnick et al., 2008), neurotransmitter release (Itakura et al., 2005), and voltage-dependent calcium channel currents (Blair and Marshall, 1997; Blair et al., 1999; Xing et al., 2006). Any of these processes could affect cone-OFF bipolar cell synaptic structure and function. An extensive series of cell specific Pappaa and IGF1R manipulations will be required to distinguish the autocrine/paracrine nature of Pappaa-IGF1 signaling and how it influences cone to OFF bipolar cell synapse formation.

## Acknowledgement

This work was supported by grants to A.H.M. (NIH T32 GM007507), H.B.H. (University of Wisconsin Hilldale Undergraduate Research Award), M.I.B. (NIH R01 GM116916), B.D.P. (NIH R01 EY017037, the Doris and Jules Stein Professorship from Research to Prevent Blindness, and an NIH Core Center Grant P30 EY025585), and M.A.W (Greater Milwaukee Foundation Shaw Scientist Award 133-AAA2656). The authors would like to thank Dr. Michael Taylor for technical advice on the electroretinograms, Drew Roenneburg (University of Wisconsin Department of Surgery Histology Core) and Ben August (University of Wisconsin Electron Microscopy Core) for technical assistance, and Dr. Cunming Duan for the *hsp70:dnIGF1Ra-EGFP* fish line.

